# A comparative analysis of liver tissue and novel primary organoid cultures from ruminants reveals species-specific immune architecture and metabolic specialization

**DOI:** 10.64898/2026.04.01.715896

**Authors:** M.E. Garner, D. Price, P. McCarron, D.J. Bartley, M.N. Faber, B. Quinn, M.W. Robinson, D Smith

## Abstract

The liver is widely considered to be one of the most conserved organs amongst vertebrates, with it being involved in blood detoxification, bile production and the metabolism of xenobiotic compounds. Liver organoids have previously been derived from several species and used as models of drug metabolism, toxicity, and fundamental tissue biology. To date, however, these models have not been developed from ruminant species, specifically cattle and sheep. Here we present the first report of the development and comprehensive characterisation of bovine and ovine liver organoids derived from primary liver tissue. When initially established, organoids from both species were comprised of KRT19- and KRT18-positive cholangiocytes. The capacity for organoids to differentiate into hepatocyte-enriched cultures was evaluated and it was noted that there was an increase in hepatocyte markers in bovine cultures. A comparative analysis of the liver tissue and organoids of both species revealed species-specific differences in gene expression, which were conserved within organoid cultures. Most notably, bovine liver tissue and organoids had enriched expression of genes associated with fatty acid uptake and storage whereas ovine samples had higher expression of genes associated with fatty acid conversion, highlighting fundamental differences between these two ruminant species. Differences in expression of cytochrome P450 family genes were identified alongside those associated with an inflammatory response specifically in bovine samples, whereas ovine samples had higher expression of genes associated with a protective immune response. Despite this, transcriptomic analysis of organoids from both species, cultured in both growth and differentiation media, revealed preserved expression of genes associated with key liver functions, including gluconeogenesis and xenobiotic metabolism. Transcripts associated with the flavin-containing monooxygenases (FMO) family were expressed in both organoid growth media and organoid development media (OGM and ODM respectively), and both species could metabolise triclabendazole into its primary metabolite triclabendazole sulfoxide, therefore validating the potential of the organoids to be applied as *in vitro* models of metabolism and/or toxicity. Overall, this study provides novel insights into differences in liver composition and function between ruminant species, as well as providing novel experimental models of the liver for both cattle and sheep.

## Introduction

The liver is responsible for the widespread detoxification of blood, production of bile to aid in digestion, and metabolism of xenobiotics (e.g. drug compounds, toxins or contaminants) and nutrients (1). Liver tissue is predominantly composed of hepatocytes, which comprise approximately 80% of the overall mass of the parenchyma. Hepatocytes are responsible for the production of metabolic enzymes and contribute towards maintaining homeostasis within the tissue (2). Running throughout the liver parenchyma is a system of intra-hepatic ducts comprised of cholangiocytes which are responsible for the final refinement and transportation of bile to the bile ducts (3). These epithelial cells line the intra-hepatic and bile ducts and express markers such as cytokeratins, EPCAM, and SOX9 (4). Although these two cell types comprise most of the mass of the liver, and are responsible for the majority of its function, other cells such as sinusoidal endothelial cells, stellate cells and tissue resident macrophages, known as Kupffer cells, are also present and contribute towards the overall structure and function of the liver (5–7). The liver is widely considered to be conserved between all mammals, showing strong similarities in the organisation of hepatic lobes, cellular composition, and overall function (8).

Livestock, including ruminants, are prone to several metabolic diseases of the liver including hepatic steatosis or fatty liver diseases, ketosis within dairy cattle, alongside other imbalances which result in reduced metabolic function and overall productivity. The resulting production losses, when combined with the impact on overall animal welfare, creates a significant burden to the agricultural industry (9,10). Alongside metabolic disorders, livestock are also susceptible to several pathogenic viral, bacterial, or parasitic infections within the liver that similarly impact animal health and welfare and cost the livestock industry billions of US dollars annually (11–16). Despite this, there are no suitable models for the *in vitro* study of the ruminant liver. There is a current lack of fundamental knowledge regarding the mechanisms of hepatic repair, host-pathogen interactions, and metabolic processes within ruminants.

Organoids are complex, stem or progenitor cell-derived, three-dimensional (3D) cell culture systems which can more accurately represent the specific microenvironment of a given tissue when compared to two-dimensional (2D) cell lines (17). Both primary tissue and pluripotent stem cell-derived liver organoids have been previously developed from other species, including human biopsy, mice, dogs, cats, and pigs, as well as reptiles (18–24). Organoids derived from primary tissue leverage the differentiation potential of oval cells, the progenitor cells of the liver located within the canal of herring, which have the capacity for bipotent differentiation into either functional hepatocytes or cholangiocytes This differentiation potential of oval cells contributes towards the regenerative capabilities of the liver following injury, through processes driven by the activation of inflammatory cytokine cascades (8,25,26).

When oval cells are cultured in media containing the growth factors WNT3A, FGF10, and HGF, as well as small molecules such as forskolin and a TGF-β inhibitor, the resulting organoids form spherical organoid structures containing a hollow luminal space (27). These organoids are typically cholangiocyte-rich, being positive for key epithelial cell markers such as KRT18, KRT19, and EPCAM (27). The differentiation capacity of oval cells has also been previously leveraged, to differentiate organoids into functional hepatocyte-rich structures capable of drug metabolism and albumin production (28–30). Moreover, the capacity of these cultures to undergo serial passage and expansion has led to their use as a drug screening platform, as well as for in-depth studies into the cellular mechanisms involved in liver function and recovery following injury (31).

Here, we sought to adapt established protocols for generating liver organoids in other mammalian species for use in cattle and sheep thereby producing the first comprehensive development and characterisation of bovine and ovine cholangiocyte and differentiated organoids. Observed differences in organoid cultivation between the two species inspired us to perform a comparative analysis which revealed surprising differences in liver biology between these two ruminant species in tissue, which was retained in primary organoids. We also demonstrate the functional capacity of the organoids for modelling drug metabolism *in vitro*.

## Results

### Establishment of Ruminant Liver Organoids

Hepatic ductal fragments were successfully isolated from both bovine and ovine liver tissue by enzymatic digestion and when seeded into Matrigel™ with organoid growth media (OGM) these fragments formed into spherical liver organoids. Small spherical structures began forming from these larger fragments within 24 hours following initial seeding. Expansion of organoids from both species occurs over several days, with the average size of a mature organoid ranging from 100-300 µm (Fig 1. B). Both bovine and ovine liver organoids could be maintained for 5-7 days at each passage, prior to further passaging by mechanical disruption. Following mechanical disruption and re-seeding into Matrigel™, fragments of organoids rapidly recover to form closed structures with a hollow lumen, resembling a liver organoid (Fig 1. C). Ovine organoids can be maintained over multiple serial passages (up to at least P20) with no change to phenotype or ability to recover observed. Unlike ovine organoids, however, bovine organoids could not be consistently maintained and propagated in culture over serial passage in expansion media. Although bovine organoids occasionally reached P5 or P6, most attempts at maintaining these cultures resulted in cells or cell fragments crashing (failure of organoids to reform following fragmentation during passaging) at approximately P3 or P4. This crashing of the bovine cultures occurred in organoids derived from animals from two separate cohorts, with organoids becoming smaller in size and darker in appearance. Nonetheless, prior to P3, bovine organoids could be passaged and expanded in number, with domes routinely observed containing at least 50-100 organoids. Both bovine and ovine liver organoids were capable of cryopreservation using Cryostor CS10 and subsequent resuscitation following storage at -150 °C (minimum 12 months) (Fig 1. D). During resuscitation, organoids were seeded into two 50 µl Matrigel™ domes to increase cellular density and cell-to-cell interactions and thereby encourage recovery. As bovine organoids could not typically be cultured past P4, cryopreservation of these organoids occurred only at P0 or P1.

**Figure 1.**
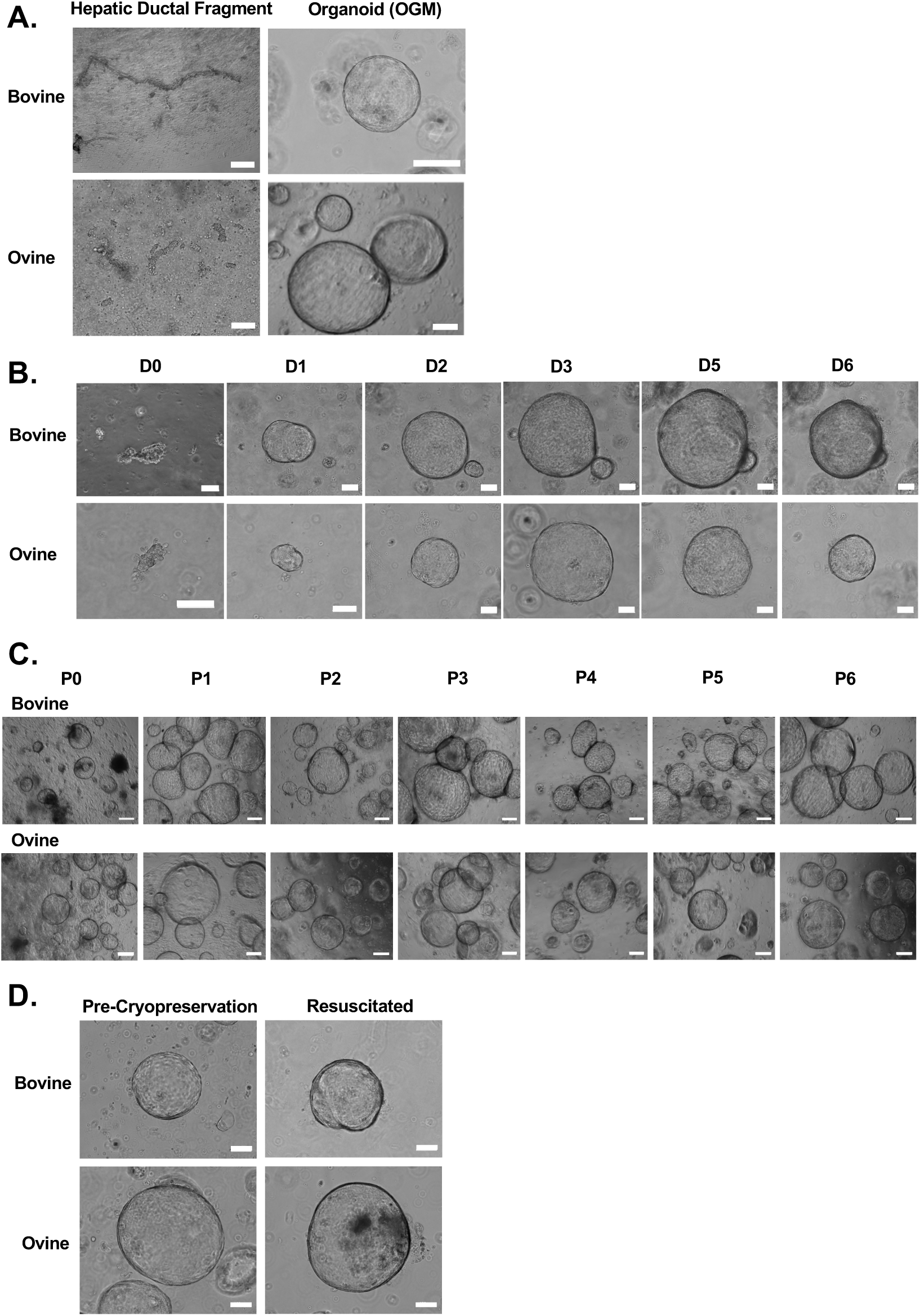
Bovine and ovine liver organoids formed from hepatic ductal fragments isolated from primary liver tissue. (A) Hepatic ductal fragments (left panels) representative of individual organoids seven days post-seeding in OGM (right panels) for bovine (top) and ovine (bottom). (B) Bovine (top) and ovine (bottom) organoid fragments at day 0 (D0) rapidly forming closed three-dimensional structures within 1-2 days and expanding in size over time to day 6 (D6). (C) Representative images of bovine (top) and ovine (bottom) organoids across serial passage (P1 – P6), up to passage 6. (D) Representative images of bovine (top) and ovine (bottom) organoids pre-cryopreservation and after being resuscitated. All images were taken using phase contrast imaging. Scale bars = 50 µm.

### Characterization of Cholangiocyte-Rich Organoids

RNA sequencing of organoids and liver tissue derived from the same individual animals was performed for a comprehensive characterisation of the cell types and states present in the organoid cultures. Principal component analysis (PCA) of organoids grown in OGM compared to tissue revealed that differences primarily lie between the tissue and organoid cultures. Importantly regarding sample reproducibility, within-group animal-to-animal variation was low for organoids and for tissue, for both species (Fig. 2.A and 2.B). Moreover, little variation was also observed at a transcriptional level between passage 2 and passage 7 organoids, indicating little variation after multiple rounds of passaging (Fig. 2. B). As bovine liver tissue and organoids were derived from two cohorts of animals of different ages, the clustering of organoids and of tissue reveals limited variation regardless of these background factors, demonstrative of good reproducibility from one animal and cohort to the next (Fig 2. A).

**Figure 2.**
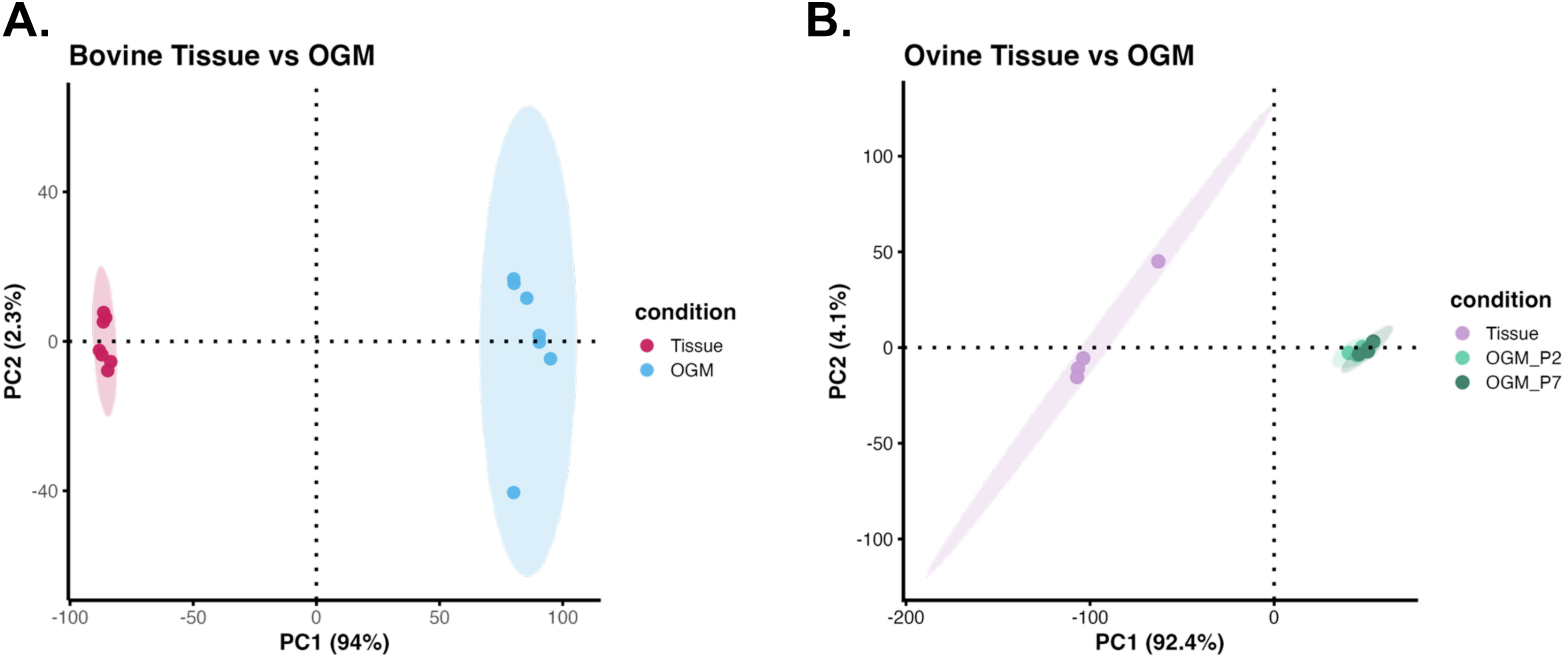
Principle component analysis (PCA) of bovine and ovine liver tissue and organoids grown in OGM display low animal-to-animal and passage-to-passage variation. (A) PCA plot of sample transcriptomes for bovine tissue and organoids grown in organoid growth media (OGM), derived from two independent cohorts (P1-2) (B) PCA plot of sample transcriptomes for ovine tissue and organoids grown in OGM, including organoids at passage 2 and passage 7.

Cluster marker enrichment analysis was performed for both species, comparing differentially expressed genes (DEGs) associated with either liver tissue or organoids. Firstly, leveraging published single cell human liver atlas data, transcript markers associated with clusters 4, 7, 24 and 39 from Aizarain et al (2019) were found to be enriched in the DEGs associated with organoids grown in OGM (Figure 3A, 4A). These clusters have previously been annotated as cholangiocytes, indicating that the bovine and ovine organoids grown in OGM are enriched for cholangiocyte transcript markers. A full list of the genes associated with each cluster can be found in Supplemental File 1, and complete spreadsheets of DEGs associated with bovine and ovine samples can be found in Supplemental Files 2 and 3, respectively.

**Figure 3.**
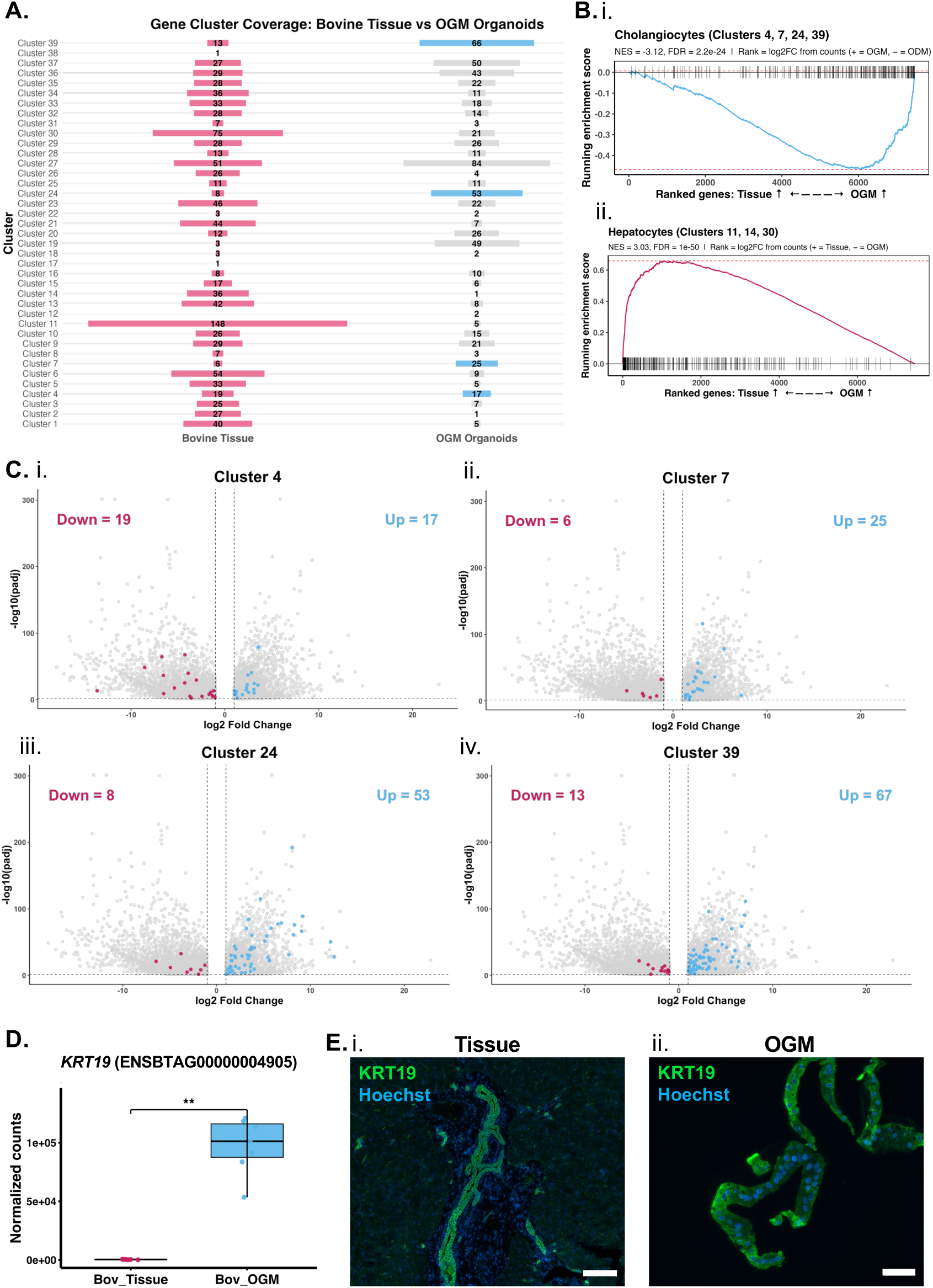
Bovine organoids are comprised of progenitor-like cholangiocytes when grown in OGM. (A) Cluster marker enrichment analysis leveraging a single cell sequencing-based atlas of the human liver (32) was applied to determine the number of cluster-enriched genes differentially enriched in bovine liver tissue and organoids grown in OGM. Transcripts associated with cholangiocyte clusters (4, 7, 24, 39) are enriched in OGM when compared to tissue. (B) Gene set enrichment analysis (GSEA) reveals transcripts associated with cholangiocyte clusters are enriched in OGM (B. i), whereas transcripts associated with the hepatocyte clusters (clusters 11, 14, 30) are enriched in bovine tissue (B. ii). (C) Volcano plots showing the differential expression of cholangiocyte cluster-enriched transcript markers in bovine tissue and OGM-cultured organoids. (D) Gene expression of the cholangiocyte and epithelial cell-associated marker *KRT19* in bovine liver tissue and OGM-cultured organoids (** *p* = 0.0022). (E) Immunohistochemistry analysis with a mouse anti-KRT19 antibody on bovine tissue shows KRT19+ expression (green) specifically of cells lining the intra-hepatic ducts within the tissue (E. i). All cells within bovine OGM-cultured organoids were KRT19+ cholangiocytes (E. ii). Scale bar represents 50 µm (E. i-ii).

**Figure 4.**
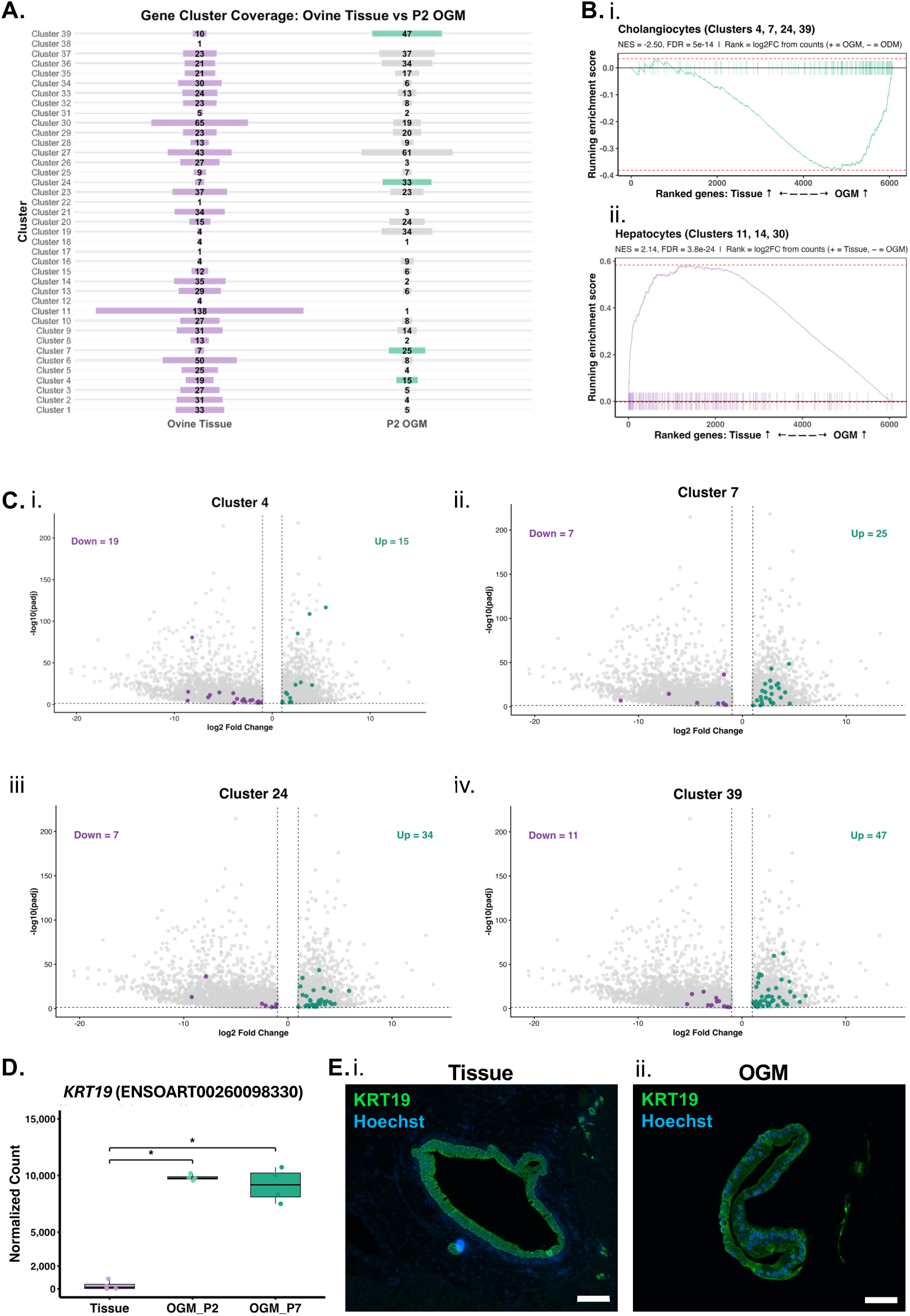
Ovine organoids are comprised of progenitor-like cholangiocytes when grown in OGM. (A) Cluster marker enrichment analysis leveraging a single cell sequencing based atlas of the human liver (32) was applied to determine the number of cluster-enriched transcripts differentially enriched in ovine liver tissue and organoids grown in OGM. Transcripts associated with cholangiocyte clusters (4, 7, 24, 39) are enriched in OGM when compared to tissue. (B) Gene set enrichment analysis (GSEA) reveals transcripts associated for cholangiocyte clusters are enriched in OGM (B. i), whereas transcripts associated with the hepatocyte clusters (clusters 11, 14, 30) are enriched in ovine tissue (B. ii). (C) Volcano plots showing the differential expression of cholangiocyte cluster-enriched transcript markers in ovine tissue and OGM-cultured organoids. (D) Gene expression of the cholangiocyte and epithelial cell-associated marker *KRT19* in ovine liver tissue and OGM-cultured organoids (** p = 0.0022). (E) Immunohistochemistry analysis with a mouse anti-KRT19 antibody on ovine tissue shows KRT19+ expression (green) specifically of cells lining the intra-hepatic ducts within the tissue (E. i). All cells within ovine OGM-cultured organoids were KRT19+ cholangiocytes (E. ii). Scale bar represents 50 µm (E. i-ii).

To further indicate the cell type specificity of OGM-cultured organoids, gene set enrichment analysis (GSEA) was performed using transcript markers associated with either all of the annotated cholangiocyte clusters (4, 7, 24 and 39) or hepatocyte clusters (11, 14, 30) in Aizarin et al (2019) (Supplemental Fig. 1; Supplemental Fig. 2). This analysis determined that transcripts associated with cholangiocytes were enriched in organoids (OGM) compared to tissue, while those associated with hepatocytes were enriched in tissue, compared to organoids (OGM). A closer analysis of all the markers within each of the cholangiocyte clusters 4, 7, 24 and 39 (Aizarain et al, 2019) is shown in the volcano plots in Figures 3C and 4C, which reveals some cholangiocyte markers were variably enriched between tissue and organoids, although the majority are enriched in organoids.

Cytokeratin 19 is recognized as a classic marker of mammalian cholangiocyte epithelial cells which comprise the intrahepatic ducts (33). RNA sequencing data revealed the increased expression of *KRT19* in both bovine and ovine organoids in growth media conditions when compared to whole tissue, highlighting the enrichment of *KRT19* expressing cells in the organoids. For ovine organoids, this enrichment of *KRT19* remained consistent between early (P2) and later (P7) passage (Fig 3. D). This was further supported at the protein level whereby KRT19-specific antibody was shown to specifically label cells lining the intra-hepatic ducts within the liver tissue from both species (Fig 3. E.i; Fig 4. E.i)). IHC analysis of both bovine and ovine organoid sections showed that every cell of the organoid (OGM) cultures labelled with the KRT19 antibody, akin to the cells lining the intra-hepatic ducts (Fig 3. E.ii; Fig 4. E.ii).

### Differentiation of Bovine and Ovine Liver Organoids

To assess the ability of ovine and bovine organoids to differentiate into more hepatocyte-like, metabolically active cells, organoid cultures were grown in OGM for three days prior to switching to Hepaticult Organoid Differentiation Media (ODM; StemCell Technologies) and grown for a further five days. Differentiated bovine organoids underwent a morphological phenotypic change, becoming darker and denser in appearance, as well reducing in size when compared to organoids in OGM (Fig 5. A). This physical change occurred approximately 72 hours after the media was switched to ODM. Alongside physical changes, RNA sequencing analysis of OGM compared to ODM for bovine organoids revealed 1646 total DEGs between the conditions (983 Up in OGM; 663 Up in ODM: Fig 5. B) (Supplemental File 2). Analysis of the genes *ALB, MKI67,* and *KRT18* revealed expression patterns associated with differentiation into hepatocyte-like cells. *ALB* expression was found to be present in both bovine and ovine tissue (Supplemental Fig. 3) and significantly upregulated (*p* = 0.0022) in ODM when compared to OGM, with transcript read count data indicating that *ALB* expression is switched off in OGM (Fig 5. C). IHC analysis using a rabbit anti-bovine serum albumin (BSA) antibody revealed that albumin was not expressed by cells in OGM (Fig 5. D.i), however upon differentiation in ODM albumin-specific labelling was observed across all cells of the organoids (Fig 5. D.ii). *MKI67* was selected as a proliferating cell marker, therefore indicative of the ability for cells within the organoids to continue to expand. When cultured in ODM, *MKI67* expression was found to be significantly downregulated when compared to OGM (Fig. 5. E). Moreover, IHC analysis with a rabbit anti-Ki67 antibody revealed that in OGM, a larger number of the cells were Ki67^+^ (Fig 5. F.i). Upon differentiation Ki67^+^ cells reduced in numbers, consistent with fewer *MKI67* transcripts in ODM (Fig 5. F.ii). This finding is consistent with these organoids undergoing terminal differentiation upon switching to ODM. *KRT18* was also found to be significantly downregulated in ODM when compared to OGM for bovine organoids, however overall transcription of this gene was not switched off in ODM (Fig 5. G). IHC analysis using a mouse anti-KRT18 antibody revealed protein expression of KRT18 in most cells of the organoid cultures in both OGM and ODM, however upon differentiation localization of KRT18 in the cytoplasm is lost and only remains at the cell membrane (Fig 5. H. i-ii).

**Figure 5.**
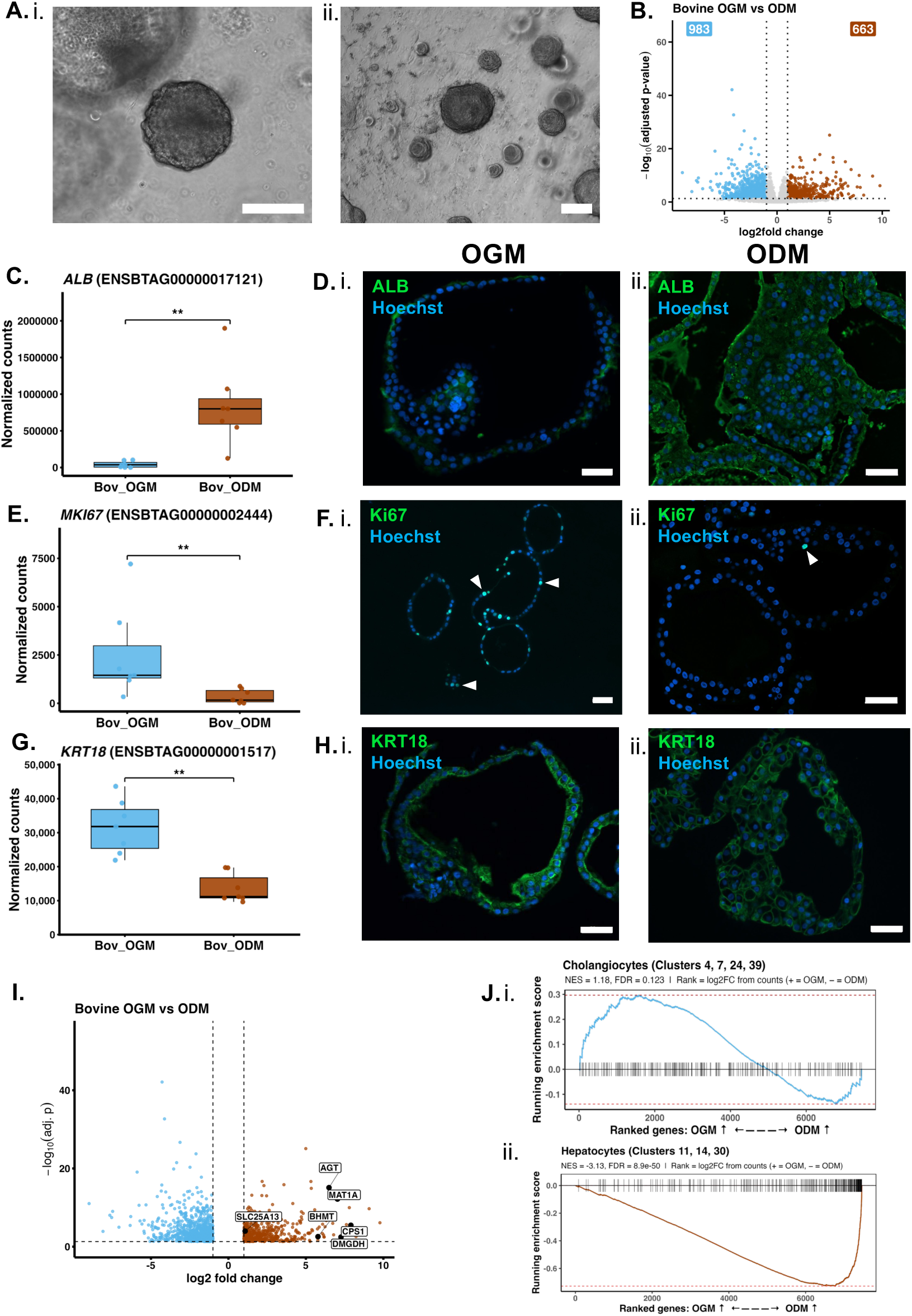
Bovine organoids undergo morphological and transcriptional changes when cultured in organoid differentiation media (ODM). (A) Bovine organoids undergo phenotypic changes, becoming darker and smaller when cultured in ODM. (B) A total of 1,646 differentially expressed genes were identified when comparing bovine organoids cultured in ODM to those in OGM. (C) Organoids in ODM had significantly higher expression of *ALB* (p = 0.0022). (D) IHC analysis revealed minimal labelling of ALB (green) in OGM (D. i) and strong labelling of most cells in ODM (D. ii). (E) Expression of the replicating cell marker *MKI67* was significantly decreased in ODM (p = 0.0073), which is also represented by a decrease in Ki67+ cells (green, indicated with white arrows) labelled in ODM (F. ii) when compared to OGM (F. i). The expression of *KRT18* was also significantly decreased in ODM (p = 0.002), however IHC indicated KRT18 (green) is lost in the cytoplasm and limited to the cell membrane in ODM (H. i-ii). (I) Analysis of genes associated with the urea cycle showed expression of 6 genes was enriched in ODM. (J) GSEA revealed that transcripts associated with cholangiocyte clusters (4, 7, 24, 39) were enriched in OGM (J. i) whereas those associated with hepatocyte clusters (11, 14, 30) were enriched in ODM (J. ii). Scale bar = 50 µm.

Alongside the increase in albumin production in bovine differentiated organoids, genes associated with the urea cycle were also found to be upregulated in ODM conditions, suggesting that upon differentiation, organoids display increased characteristics associated with hepatocytes (Fig 5. I). GSEA was performed leveraging the clusters associated with cholangiocytes (4, 7, 24, 39) and hepatocytes (11, 14, 30) from Aizarin et al., 2019. GSEA revealed that whilst genes associated with cholangiocytes are enriched in OGM compared to ODM, there is a subset of genes from these clusters that are preferentially expressed in ODM (Fig 5. J.i; Supplemental Files 2 & 3). Despite this, however, upon differentiation the genes associated with hepatocytes become enriched, again indicating the switching of bovine organoids from OGM to ODM results in differentiation from cholangiocytes to hepatocyte-like cells (Fig 5. J.ii; Supplemental Files 2 & 3).

Differentiation of ovine organoids also resulted in a change of phenotype, with these organoids also becoming smaller and darker in appearance (Fig 6. A). DEG analysis identified a total of 1903 DEGs (1206 Up in OGM; 697 Up in ODM) when comparing ovine organoids grown in ODM to those grown in OGM (Fig 6. B). The expression levels of *ALB, MKI67,* and *KRT18* was also investigated for differentiated ovine organoids. Unlike bovine cultures, no increase in *ALB* expression was observed upon differentiation of ovine cultures (Fig 6. C). *ALB* expression, however, was much higher in ovine OGM conditions when compared to bovine OGM (Fig 5. C; Fig 6. C). However, this higher expression was not observed at a protein level, as although IHC analysis of organoids cultured in OGM and ODM showed some albumin labelling, the intensity of this signal was lower than in bovine ODM (Fig 6. D.ii). The reactivity of the BSA antibody to ovine albumin was confirmed by Western blot (Supplemental Figure 3). Similar to bovine ODM cultured organoids, gene expression patterns for both *MKI67* and *KRT18* showed significant decrease in ODM-cultured ovine organoids compared to those in OGM (Fig 6. E; Fig 6. G). This was confirmed through IHC analysis, where Ki67+ cells were only present in organoids grown in OGM (Fig 6. F.i-ii). Alongside this, cells within the ovine organoids were all KRT18+ in OGM, whereas upon differentiation this was decreased with fewer cells positive for KRT18 and similar to bovine KRT18 localization became less cytoplasmic (Fig 6. H.i-ii).

**Figure 6.**
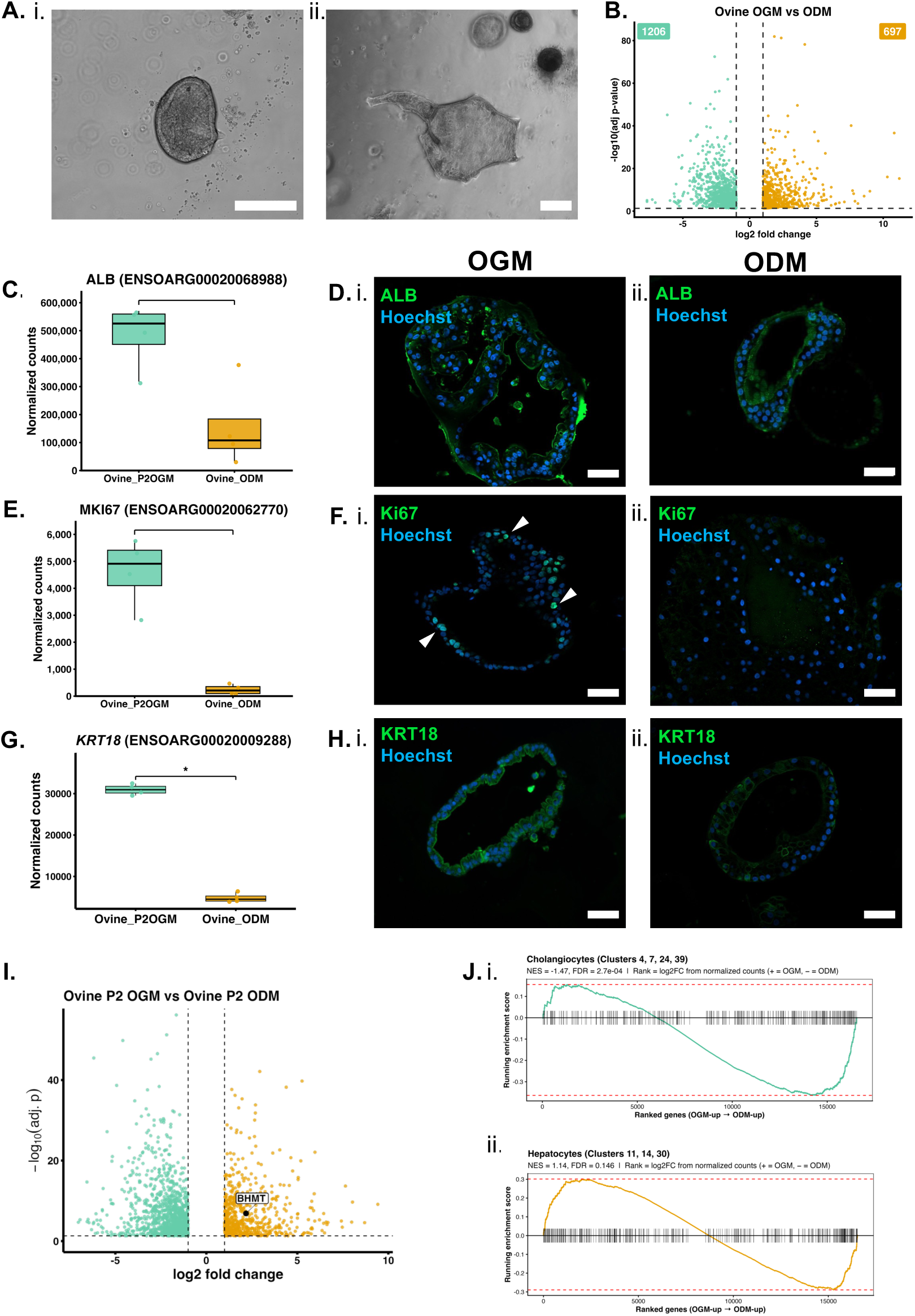
Ovine organoids cultured in ODM remain rich in a subset of cholangiocyte markers. (A) Ovine organoids become smaller, denser, and more irregularly shaped when cultured in ODM. (B) A total of 1,903 DEGs were identified in ODM when compared to ODM. (C) *ALB* expression is well maintained in OGM and ODM (p = 0.08), with labelling of ALB (green) remaining consistent through IHC analysis (D. i-ii). The expression of key genes *MKI67* (E) and *KRT18* (G) both were significantly decreased in ODM (p = 0.03 and p = 0.0304 respectively), with this decrease in gene expression also being translated to a decrease at a protein level through IHC (green) labelling with anti-Ki67 (F. i-ii) and anti-KRT18 antibodies (H. i-ii). White arrows in panel F.i. indicate Ki67-positive nuclei. (I) Analysis of genes associated with the urea cycle revealed only *BHMT* was significantly increased in ODM. (J) GSEA of cholangiocyte-associated clusters (4, 7, 24, 39) identified enrichment of these clusters in both OGM and ODM (J. I), whereas hepatocyte-associated clusters (11, 14, 30) were more enriched in OGM (J. ii). Scale bar = 50 µm.

Cytokeratin 19 (KRT19) expression followed a similar pattern to KRT18 whereby although *KRT19* transcription persisted in ODM-cultured organoids, there was a significant decrease in *KRT19* in both bovine and ovine ODM compared to OGM-cultured organoids. Moreover, the localization pattern of KRT19 in ODM-cultured organoids became less cytoplasmic, predominantly presenting like rings demarcating individual cell borders in KRT19 antibody-labelled sections (Supplemental Fig. 4).

The same set of genes associated with the urea cycle were explored for enrichment in ovine differentiated cultures, and although all six genes were expressed in both OGM and ODM, only *BHMT* was found to be significantly upregulated upon differentiation (Fig 6. I; Supplemental Files 2 & 3). However, the other genes are well-expressed in OGM (unlike in bovine OGM where transcript counts are low or absent for these urea cycle markers) and collectively all urea cycle markers are consistently more highly expressed across ovine organoid samples compared to bovine, regardless of media conditions (Supplemental Files 2 & 3). This indication of limited hepatocyte differentiation in ovine ODM organoids was supported by GSEA analysis on transcript markers in clusters associated with cholangiocytes, which revealed that there was a similar number of genes in these clusters that were enriched in OGM and ODM (Fig 6. J.i; Supplemental Files 2 & 3). Alongside this, the genes associated with the hepatocyte clusters were enriched in OGM when compared to ODM, consistent with high *ALB* expression in OGM-cultured ovine organoids. Collectively, this suggests that ovine organoids cultured under growth conditions partially exhibit hepatocyte properties, but do not undergo a complete cholangiocyte to hepatocyte differentiation to the same extent as bovine organoids when cultured in differentiation media (Fig 6. J.ii; Supplemental Files 2 & 3).

### Investigation of Species-Specific Differences

The observed differences between bovine and ovine cultures reported thus far encouraged us to investigate species differences further. Firstly, all conditions from each species (including tissue) were compared using PCA. While separation along PC1 can be identified as tissue vs organoids (regardless of species), PC2 can clearly be identified as sample separation based on species (Fig 7. A). Collectively, this suggests large differences in the transcriptome of the bovine liver compared to the ovine liver, and that tissue-derived organoids retain a transcriptomic signature associated with the specific host species they are derived from. This is supported by comprehensive differential expression analysis which revealed a surprisingly high number of DEGs (4824) between bovine and ovine tissue (Fig 7. B.i). Similar numbers of DEGs were also identified between ovine and bovine OGM organoids (5077; Fig 7. B.ii), and between ovine and bovine differentiated organoids (5223; Fig 7. B.iii; Supplemental File 4). Of these observed differences, 853 DEGs were conserved between all conditions for bovine samples (Fig 7 C.i) and 891 for ovine samples (Fig 7. C.ii), which therefore represent unique transcriptomic signatures of each individual species that are retained within the organoid cultures. Gene ontology (GO) term analysis revealed terms such as positive regulation of T cell proliferation, regulation of leukocyte mediated cytotoxicity, ribosomal subunit, and respiratory chain complex IV for genes associated with the bovine samples (Supplemental Fig. 5). GO term analysis of genes shared between ovine samples revealed terms such as translation, cytoplasm, structural molecule activity, and rRNA binding (Supplemental Fig. 5).

**Figure 7.**
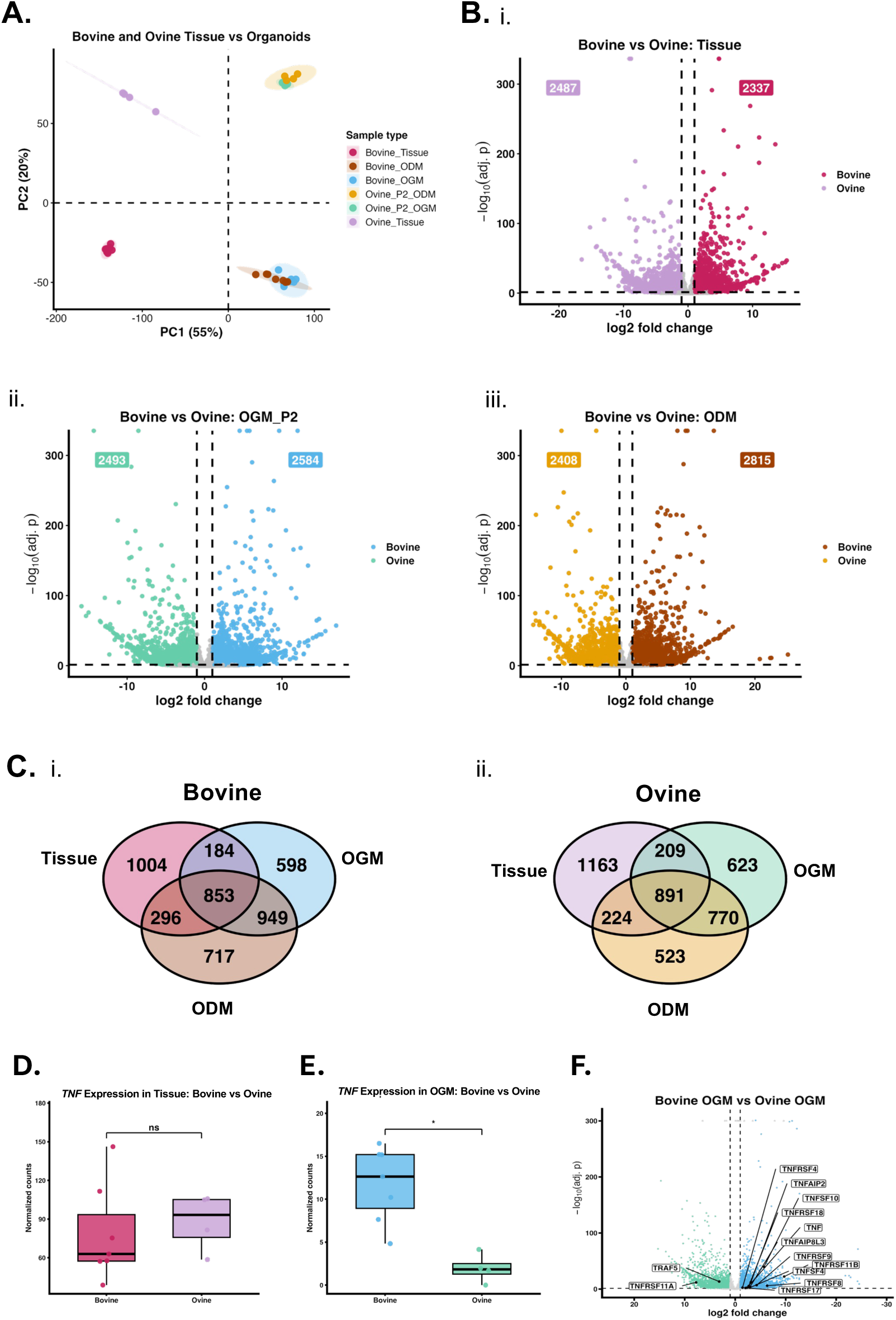
Species-specific differences are conserved between tissue and organoids grown in both OGM and ODM. PCA analysis demonstrates that organoids retain species-specific characteristics with PC1 separation based upon tissue or culture, whereas PC2 is based upon species (A). When comparing bovine and ovine tissue (B. i), organoids in OGM (B. ii) or organoids in ODM (B. iii), there are approximately 5000 total DEGs identified between the species in each condition. Of these DEGs, 853 (C. i) and 891 (C. ii) differences were retained in all three conditions for bovine and ovine samples respectively. To further investigate the differences in differentiation between bovine and ovine organoids, expression of *TNF* was compared between (D) tissue and (E) organoids in OGM, noting equal expression of *TNF* in tissue and significantly enriched expression of *TNF* in bovine organoids cultured in OGM (p < 0.05). (F) A total of 11 TNF-associated genes were found to be enriched in bovine OGM when compared to ovine OGM, both at P2.

As previously described, TNF-α has been linked to the differentiation and proliferation of hepatocytes following liver injury, with differentiation media conditions often containing TNF-α to drive the cholangiocyte to hepatocyte differentiation. It was observed that supplementation of TNF-α in ODM for organoid differentiation would induce rapid cell death in bovine cultures. Alongside this, the bovine liver organoids developed here are only stable in OGM culture up to approximately passage four, at which point they typically fail to passage further. This therefore provided a clue that endogenous TNF-α levels might already be elevated in bovine cultures and tissue and that this could underpin the inability to serially passage them in OGM. Therefore, the expression of *TNF* and *TNF*-associated genes in bovine and ovine tissue, as well as organoids in growth media, were compared. No significant difference was observed between the *TNF* expression levels in tissue (Fig 7. D), however *TNF* was significantly upregulated (p = 0.0304) in bovine OGM when compared to ovine organoids grown in OGM (Fig 7. E). Upon further investigation, it was found that 11 *TNF*-associated genes were upregulated in bovine cultures (Fig 7. F).

Following this observation, we sought to leverage this information to improve the hepatocyte differentiation of ovine liver organoids. A series of differentiation media conditions were tested using varying concentrations of TNF-α supplementation, alongside the presence or absence of a TGF-β inhibitor (A83.01). Organoids were grown in culture for 3 days in standard OGM, prior to switching to the different ODM conditions for a further 5 days. Following RNA extraction and cDNA synthesis from each of these conditions, RT-qPCR was performed to evaluate the expression of *ALB* across the different conditions. *ALB* expression remained consistent across most ODM conditions, with relative expression levels remaining similar regardless of the addition or removal of TNF-α or A83.01 (Supplemental Fig. 6). To investigate whether the change in media conditions altered the amount of ALB protein present in the cells, fixed sections of organoids grown in OGM, ODM, and ODM with increased supplemental TNF-α and no A83.01 were labelled using an anti-BSA antibody. No difference in the ALB labelling or overall fluorescence intensity was observed between ovine organoids grown in standard ODM or ODM containing increased TNF-α and no A83.01 (Supplemental Fig. 6).

Aside from TNF being elevated in bovine organoids, GO terms associated with other immunological markers and responses were enriched in the bovine samples. To further explore these innate differences in the immune landscape between the species, we next identified all genes within the species-specific DEGs that were related to the immune response and found a subset of genes that were also differentially enriched between bovine and ovine samples (Fig 8. A) (Supplemental File 4). In total 15 genes were found to be specifically enriched in bovine samples and 10 genes enriched in ovine samples. Of these, the genes enriched in bovine tissue and organoids were found to be associated with a more inflammatory response, with enrichment of *IL-1β, VSIG4* and *LTB4R2* (34–38). The expression of *VSIG4*, a Kupffer cell-specific marker, within the bovine samples indicates that there may be immune cells present within the organoid cultures, however this was not observed through microscopy-based analysis. Those genes enriched in ovine samples, however, were more associated with a protective response, such as *ICOS, IFI6, IL-17RE* and *EOLA1* (39–47).

**Figure 8.**
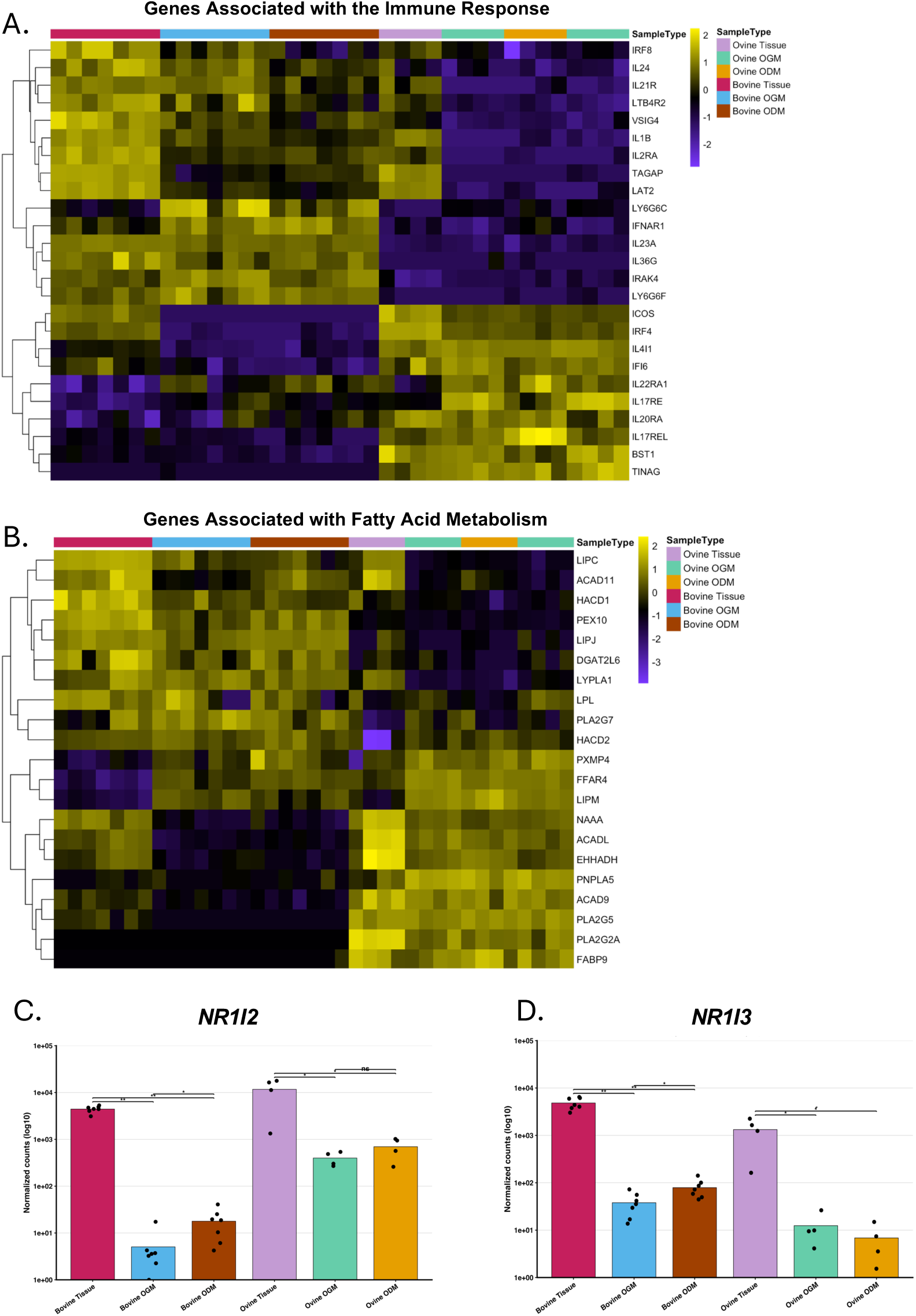
Species comparison reveals differences in xenobiotic and fatty acid metabolism, as well as differences in the immune landscape. (A) Species-specific differences observed in immune response genes were observed with 15 genes specifically enriched in bovine samples and 10 genes enriched in ovine samples. (B) For further comparison between the species, genes associated with fatty acid metabolism were evaluated, with 8 genes specifically enriched in all bovine samples and 14 specifically enriched in ovine samples. Next, genes involved in xenobiotic metabolism were investigated and it was found that (C) *NR1I2*, encoding the pregnane X receptor (PXR) xenobiotic metabolism pathway receptor, was specifically enriched in ovine samples, (D) whereas *NR1I3,* which encodes the constitutive androstane receptor (CAR) pathway receptor, was specifically enriched in bovine samples.

Another key difference between cattle and sheep is bovine susceptibility to hepatic lipidosis, or fatty liver disease (48–50). Using a similar approach as with the immunological markers, we then identified a subset of genes associated with the metabolism of fatty acids that were preferentially enriched in a species-dependent manner. We identified 8 genes associated with this process that were specifically enriched in bovine samples, and 14 genes that were enriched in ovine samples (Fig 8. B). Those enriched in bovine tissue and organoids were associated with lipid uptake, storage, and processing, whereas those associated with the ovine samples are involved in fatty acid β-oxidation and lipid remodelling (51,51–55) (Supplemental File 4).

As markers for fatty acid metabolism were differentially enriched between the species, we then looked at other markers of metabolic processes. The genes *NR1I2* and *NR1I3*, which are integral to the process of xenobiotic metabolism, were found to be differentially enriched between the species (Fig 8. C, D). *NR1I2*, also referred to as *PXR*, was found to be enriched in all ovine samples, whereas the alternative xenobiotic metabolism pathway *NR1I3*, or *CAR*, was found to be specifically enriched in bovine samples (56,57). Cytochrome (CYP) P450 enzymes are associated with drug, lipid, and vitamin metabolism alongside the alteration of bile for secretion (58–61). As such, these enzymes are associated with expression in hepatocytes. We evaluated the expression of *CYP* genes within bovine and ovine tissue and organoids and found 29 *CYP*s annotated and shared between both the bovine and ovine genomes (Fig 9. A; Supplemental Files 2 & 3). The expression patterns of some *CYP*s were shared between the species, such as *CYP1A1, CYP1B1,* and *CYP51A1,* which were transcriptionally enriched in both OGM and ODM compared to tissue. A subset of CYP genes were also preferentially expressed in organoids cultures in OGM from both species, when compared to ODM such as *CYP2S1* and *CYP20A1*. Alongside this, the expression of some CYP genes were found to differ between the species, such as *CYP4B1, CYP4B8* and *CYP4X1* which were expressed in both bovine and ovine tissue, however, were only expressed in bovine organoids. Some CYP genes were also found to be specifically enriched in either bovine (*CYP2F1, CYP2S1, CYP4F8, CYP4F22, CYP4X1, CYP24A1*) or ovine (*CYP2U1, CYP2W1, CYP7B1*) organoids in OGM. Further to this, 16 CYPs were found to be uniquely annotated only within the bovine genome, including *CYP2C23, CYP2J30, CYP26C1,* and *CYP27B1* which were primarily expressed within the organoids (Fig 9. B.i). Of the 8 CYP genes uniquely annotated to the ovine genome, *CYP17A1* was expressed in both OGM and ODM, whereas *CYP2C18, CYP2J2,* and *CYP3A5* were enriched in ODM when compared to OGM (Fig 9. B.ii).

**Figure 9.**
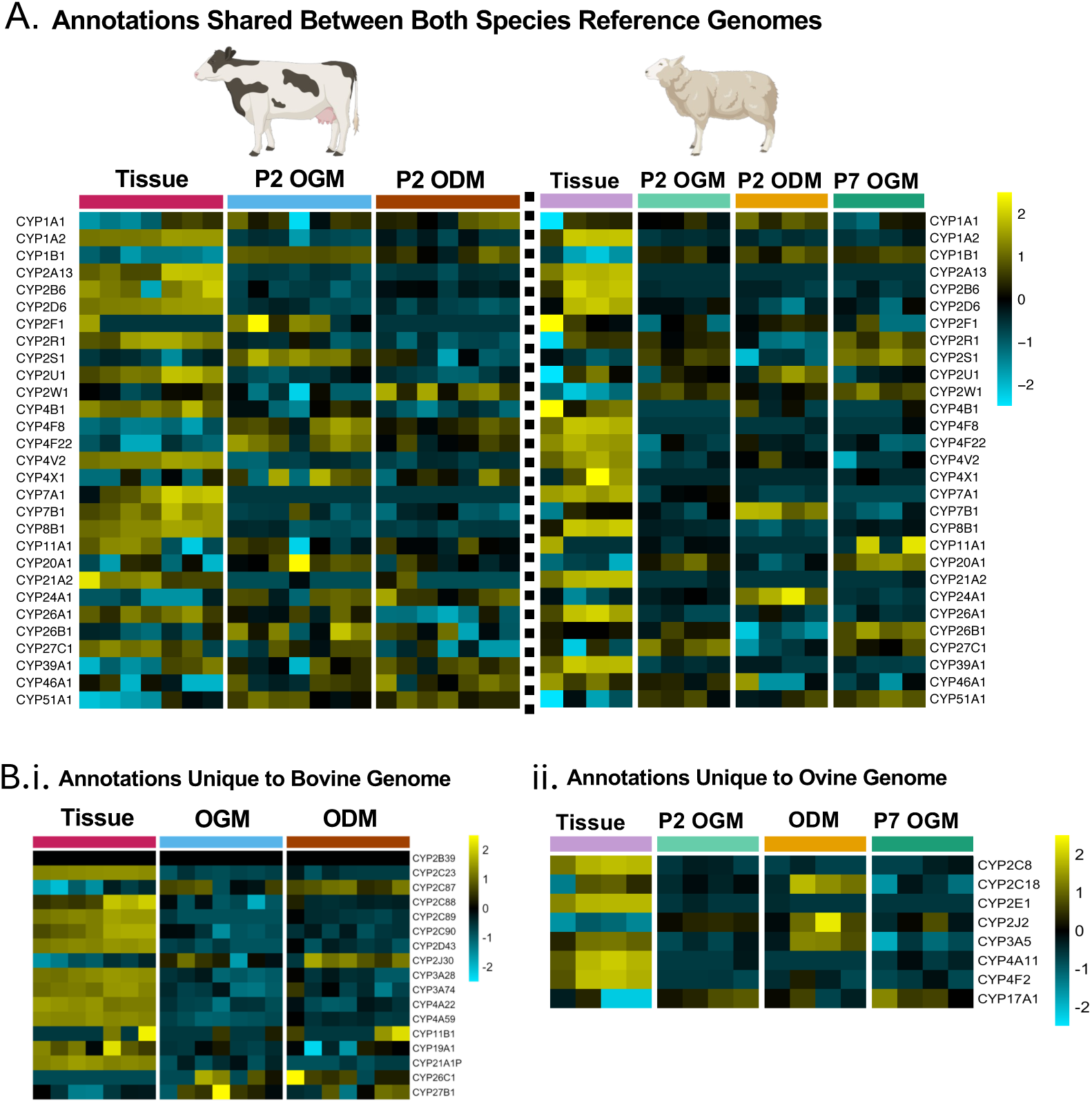
Comparison of the expression of cytochrome P450 enzymes in bovine and ovine liver tissue and organoids. (A) In both bovine and ovine tissue and organoid samples, 29 genes encoding CYP P450 enzymes were annotated in both reference genomes, with expression patterns varying depending on species and organoid condition. Apart from the 29 shared annotations, (B) 16 CYP genes were uniquely annotated in the bovine genome and (C) 9 CYP genes were uniquely annotated in the ovine reference genome. Expression patterns of tissue samples, organoids in OGM, and organoids in ODM are displayed.

Although the liver is not the sole organ responsible for the production of glucose, it is the site in which in the majority of endogenous glucose is produced through the process of gluconeogenesis in which lactate is first converted to pyruvate and subsequently to glucose (62–64) (Fig 10. A). The gene products responsible for both the lactate to pyruvate (*LDHA, LDHB)* and pyruvate to glucose conversion (*PC, PCK1, PCK2, FBP1, G6PC1*) were all expressed in both bovine (Fig 10. B) and ovine (Fig 10. C) tissue and organoids (Supplemental Files 2 & 3). All genes were found to be expressed in all conditions, with similar trends of expression in the organoids found between bovine and ovine cultures. *LDHA* was enriched in both bovine and ovine organoids when compared to tissue, whereas expression of *LDHB* and *FBP1* was at a similar level between tissue, OGM, and ODM in both species.

**Figure 10.**
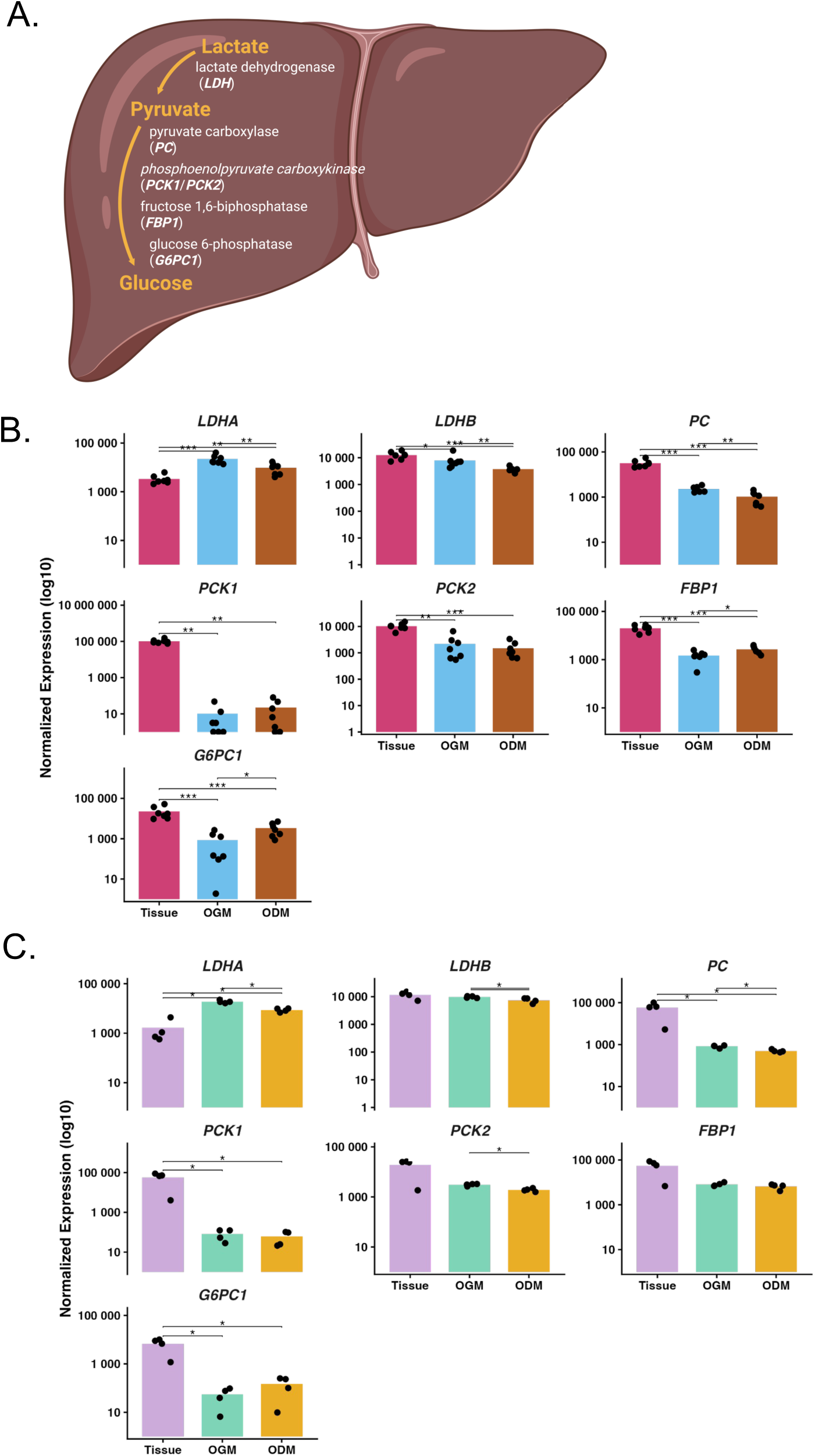
Evaluation of genes associated with gluconeogenesis in bovine and ovine organoid cultures. (A) Gluconeogenesis is the process by which lactate is first converted to pyruvate via lactate dehydrogenase (*LDHA, LDHB*) and then subsequently converted to glucose via four specific enzymes encoded by *PC, PCK1, PCK2, FBP1,* and *G6PC1.* All seven genes associated with gluconeogenesis were expressed in both bovine (B) and ovine (C) tissue and organoids. Expression of *LDHA* was significantly enriched in both bovine OGM (B) and ovine OGM (C). * = p < 0.05, ** = p < 0.01, *** = p < 0.001. Figure (A) was generated using BioRender.com.

### Ruminant Liver Organoids as an *in vitro* Model of Drug Metabolism

To evalute the potential for the developed ruminant liver organoids to be used as functional *in vitro* models of liver drug metabolism, the expression of genes within the flavin-containing monooxygenase (FMO) family were explored for both species, based on their role in the detoxification and metabolism of drug compounds. All five genes of the FMO family were expressed in both bovine (Fig 11. A) and ovine (Fig 11. B) tissue. The expression level of *FMO4* was similar between both species across tissue and both organoid conditions (OGM and ODM). Bovine organoids cultured in ODM had increased expression of all FMO genes, *FMO1-5,* when compared to OGM. This was not observed in ovine organoids, with the gene expression levels of both *FMO1, FMO2, FMO3,* and *FMO5* significantly lower than observed in tissue (Fig 11. B; p < 0.05). Expression of *FMO4* in ovine samples was similar between tissue, OGM, and ODM (Fig 11. B). To determine whether both bovine and ovine liver organoid cultures are functionally capable of drug metabolism, organoids from each species cultured in either OGM or ODM were treated with 20 µM triclabendazole (TCBZ) for 24 hours (65–67). The parent compound TCBZ was detectable by liquid chromotography/mass-spectrometry (LC/MS) within the organoids from both species cultured in both OGM and ODM (Fig 11. C; Fig 11. D). The primary TCBZ metabolite and active form, TCBZ-sulphoxide (TCBZ-SO), was also detectable in both the organoids (Fig 11. C; Fig 11. D) and the culture supernatants from both species (Fig 11. E; Fig 11. F). The amount of TCBZ-SO detected was higher in the supernatants than in the organoids themselves for both species. Collectively, this data indicates that the parent compound TCBZ was taken up by the organoids and metabolized intracellularly to TCBZ-SO which was subsequently released into the culture supernatants where it accumulated. Additional TCBZ metabolites, including TCBZ-sulphone (TCBZ-SO_2_) and keto-TCBZ, were not detected in either the organoid fraction or the cell culture supernatants, indicating that no further metabolism of TCBZ-SO into derivatives is performed by hepatic cells (Supplemental File 5).

**Figure 11.**
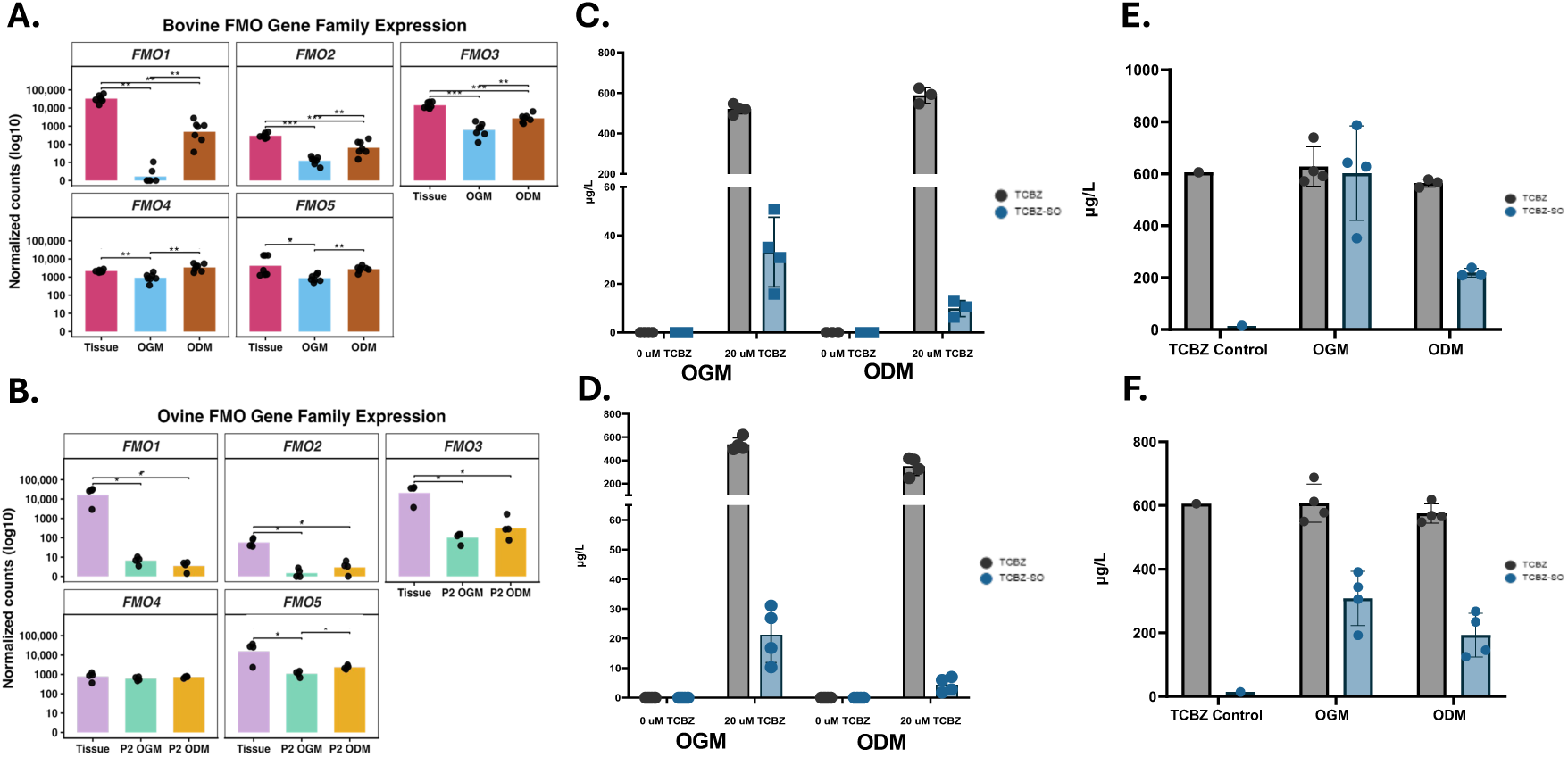
Ruminant liver organoids are metabolically functional, and actively convert triclabendazole (TCBZ) to its sulphoxide metabolite (TCBZ-SO). (A) Analysis of genes within the flavin-containing monooxygenase (FMO) family identified expression of *FMO1-5* in tissue, organoids in OGM, and organoids in ODM for both bovine (A) and ovine (B) samples. Following challenge of organoids with 20 µM TCBZ, LC-MS analysis identified both the parent drug, and its primary TCBZ-SO metabolite within the organoid fraction for bovine (C) and ovine (D). Increased TCBZ-SO was detected in organoid culture supernatants for bovine (C) and (F). Bar charts (B, C, E, F) were generated using GraphPad Prism 10 (v. 10.6.1).

## Discussion

The liver is a complex organ essential for the detoxification of blood, metabolism of drugs and nutrient compounds, and the production of bile for food digestion. Pathologies of the liver include metabolic diseases, such as hepatic lipidosis or ketosis and infectious agents including viral (e.g. Hepatitis E Virus), bacterial (e.g. *Fusobacterium necrophorum*), and parasitic (various helminth and protozoa species) infections (11,68–70). Where they occur in livestock, all these conditions impact overall animal welfare and productivity and contribute substantially towards global economic losses. While liver models have been previously developed for some species, such as porcine liver organoids for high-throughput drug testing or the use of immortalized bovine liver cell lines such as BFH12, there remains a lack of complex, physiologically relevant liver models for ruminants (71). Moreover, primary organoids have been shown to more accurately recapitulate the native microenvironment of their tissue of origin when compared with immortalized cell lines (72). Consequently, establishing a complex liver model for ruminants would represent a significant advancement for the field.

Accordingly, the present study aimed to develop a functional and physiologically relevant, *in vitro* liver model for both cattle and sheep which would be useful in a wide variety of research applications. We aimed to leverage techniques and protocols previously developed for liver organoid cultivation in other species, such as humans and mice, and apply these towards the development of ruminant liver organoids. A key feature of organoid cultivation is the baseline maintenance of a progenitor-like state, with the capacity to differentiate into other cell types upon stimulation. Previous studies indicate that liver-derived organoids are well suited for this purpose, as they typically exhibit a cholangiocyte-rich composition during early establishment and closely reflect the intra-hepatic ductal structures from which they are derived (22,27,29,30).

Following the isolation of intra-hepatic ductal fragments from primary liver tissue of both cattle and sheep, we successfully established self-organising, spherical organoid structures containing a hollow luminal space from both species. The established cultures not only phenotypically resembled liver organoids, based on their 3D architecture, but further characterisation confirmed the presence of cholangiocyte-specific markers in growth media, with each positive for KRT18 and KRT19, confirming that the OGM organoids consist of bile duct-like epithelial cells. We also showed that hepatocytes can be enriched in differentiation media, with decreased KRT18 expression and cell proliferation and increased ALB labelling (particularly bovine) in ODM organoids. Alongside this, we leveraged a single cell sequencing based atlas of the human liver (32) to perform an unbiased analysis of the cell types present in the organoids, further validating the IHC analysis. GSEA confirmed the enrichment of cholangiocyte-associated transcripts in both bovine and ovine OGM cultures and enrichment of hepatocyte-associated transcripts in ODM. This is further supported by the enrichment in ODM organoids of transcripts associated with the urea cycle, such as *CPS1*, and drug metabolism, such as those within the *FMO* family (*FMO1-5*). Interestingly, we observed a significant enrichment of both *FMO3* and *FMO5* in the differentiated organoids from both species when compared to OGM. These two genes are the primary FMO types found within the adult liver, as throughout development there is a shift from reliance on *FMO1* to *FMO3* and *FMO5* (73). OGM organoids from both species could be successfully cryopreserved and revived, demonstrating their potential to support the reduction of animal use in research in line with the 3Rs principles. Once biobanks of primary tissue-derived organoids are established, the need to return to animals for new *in vitro* cultures is substantially reduced (74).

Although cultures from both species could undergo serial passage to a greater or lesser extent, a notable difference was that bovine cultures could only be reliably maintained up to passage four. Bovine organoids were derived from two separate cohorts (spanning seven animals in total) to attempt to mitigate this issue, however this remained a consistent observation. During later passages, bovine organoids underwent phenotypic changes, whereby the luminal space became dense and dark. Cell viability in these dense organoids was compromised, leading to cell death in culture and loss of organoid-forming capacity. Intriguingly, this suggests that activating the Wnt signalling pathway in mammalian epithelial cells is not sufficient to support uncapped long-term cultivation of organoids derived from *ex vivo* progenitor cells in certain species, including the cattle-derived cultures presented here.

Lei et. al. (2025) previously reported the development of 3D bovine liver cultures. Although this study demonstrates the growth of 3D structures with a hollow luminal core in culture, there is no evidence or comment on their ability to be serially passaged. In addition, the IHC staining presented is inconsistent with localisation patterns previously reported for their chosen antibody markers, such as the tight junction marker ZO-1 and the hepatocyte marker ALB (75). The lack of commentary on the ability to serially passage these cultures (and presentation of only P1 cultures), when combined with the results presented here, suggest that the challenges in sustaining bovine culture remain unresolved and warrant further investigation.

To understand the cellular process underpinning this eventual collapse in culture viability, we compared gene expression profiles from the bovine organoids and tissue to their ovine equivalents. A surprisingly high number of DEGs were identified between the species, with approximately 1,000 of these conserved across tissue and organoid sample types. These DEGs represent the fundamental differences in the liver between these two ruminant species, in a tissue that is often considered highly conserved between all mammals. GO term analysis of these DEGs indicated that bovine samples were associated with immune-related cytotoxicity, highlighting a potential inflammatory response of the cells. In particular, *TNF and TNF-*associated gene expression was found to be higher in bovine organoids compared to ovine. TNF-α can induce differentiation (e.g. to hepatocytes), but sustained elevated levels can result in apoptosis (76), leading us to predict that endogenous TNF-α in bovine cultures eventually leads to cell death and an inability to cultivate bovine liver organoids long-term.

Among the species-conserved DEGs, we also found that bovine tissue and organoid samples were enriched in transcripts such as *IL-1B* and *VSIG4,* consistent with an elevated baseline inflammatory landscape compared to ovine (34,35). In particular, the enrichment of VSIG4, a unique marker of Kupffer cells (34) across all bovine samples might reflect an increased abundance of these tissue resident macrophage-like cell types in bovine liver tissue, leading to raised Kupffer cell carryover into organoid cultures. By contrast, ovine samples were enriched in transcripts associated with immune regulation and a protective response, such as *ICOS, IFI6,* and *EOLA1* which have previously been associated with anti-apoptotic activity, as well as repair during liver injury (39–41,43,47).

The metabolic transcriptional landscape also differed significantly between the two species, which was also reflected in their respective organoid cultures. For instance, our analysis of markers for xenobiotic metabolism and fatty acid metabolism suggested that the pathways underlying these key liver functions are fundamentally distinct between cattle and sheep, at least at a transcriptional level. Regarding xenobiotic metabolism, *NR1I2* expression was higher in bovine tissue and organoids, while *NR1I3* was higher in ovine samples, leading us to speculate that the PXR pathway could be more active in cattle, while the CAR pathway could dominate in sheep (77–79). With respect to fatty acid metabolism, we found that bovine tissue and organoids were enriched in transcripts for *LPL, LIPC,* and *DGAT2L6,* which are correlated with the uptake and storage of fatty acids within the liver, potentially highlighting a bias in bovine samples for lipid uptake rather than conversion to triglycerides (52,54,80,81). Ovine samples, on the other hand, showed higher expression of *ACADL, ACAD9,* and *ACAD11, a* subset of genes encoding acetyl-CoA dehydrogenases involved in the β-oxidation of fatty acids and their conversion to triglycerides (51,55,82). This, alongside the specific enrichment of other genes such as *EHHADH, PLA2G2A,* and *PNPLA5*, indicate that fatty acid conversion, rather than storage, is promoted in ovine liver (83–86). The bias towards fatty acid storage rather than conversion in both bovine tissue and organoids sheds light on the susceptibility of cattle to metabolic diseases, such as hepatic lipidosis (9,49,50,87). Our analysis illustrates a novel approach for comparing the transcriptional profiles of tissue and organoids from two closely related species (in this case cattle and sheep) to better understand fundamental differences in tissue physiology.

It has previously been stipulated that the bovine liver has a different cytochrome P450 landscape. For example, CYP3A4, a dominant enzyme in the liver tissue of other species (e.g. human) is not present in any annotated bovine resource (88). Here, we found that the expression profile of CYPs differed by species, with a number of *CYP* genes showing higher expression in the tissue of one species compared to the other. Moreover, we also found that some *CYP* genes are enriched in organoid cultures compared to tissue and that this could be species-specific. For example, CYPs involved in drug and xenobiotic metabolism (61,89) e.g. (*CYP1A1* or *CYP1B1*) were enriched in organoids for both species while *CYP4F8* and *CYP4F22* were only enriched in bovine organoids. Cytochrome P450 enzymes can perform a wide range of functions beyond drug metabolism, with several family members implicated in lipid and vitamin metabolism, as well as bile acid synthesis (60,90–93). *CYP7B1* has been linked to the synthesis of bile from cholesterol, and interestingly, this gene was enriched only in tissue (from both species) and ovine organoids grown in ODM (94). This observation supports our conclusion that cholangiocytes persist within the ovine organoids under differentiation conditions. A study by Bregante et. al. (2026) described the presence of either cycling cholangiocyte progenitor cells, fated towards mature cholangiocyte development, or the enrichment of bipotent progenitor cells following changes to media formulations. Interestingly, our normalized transcript count data shows bovine ODM organoids to be enriched for transcript markers associated with both hepatocytes and bipotent progenitor cells (*PROM1, FOXO3, FGFR3, HNF4A*), however these were not enriched in ovine ODM organoids. Although there was a similar level of expression for the ductal progenitor markers *SOX4* and *SOX9* in both bovine and ovine OGM and ODM, ovine organoids had higher expression of transcripts associated with mature cholangiocytes, such as *AQP1* (95). These findings highlight that although organoids from both species were derived from similar material and cultured under the same media conditions, there are observed differences in their progenitor cell state and potential to undergo hepatocyte differentiation.

When comparing all conditions between bovine and ovine samples, it was observed that 19 CYP genes were specifically annotated only in the bovine genome and 8 were restricted to the ovine genome. Although these patterns may reflect differences in the annotation of the available reference genomes, it is also possible that these genes correspond to fundamental biological differences between the bovine and ovine liver. Further exploration of these genes, alongside those annotated in both genomes but differentially expressed between species or conditions, could provide valuable insight into species-specific adaptations in liver function.

In this study, we also evaluated the expression of genes associated with gluconeogenesis, the process of lactate conversion to glucose. Although this process is not specific to the liver, as it occurs at other sites such as the kidneys and intestinal epithelia, most endogenous glucose is produced via this pathway specifically within the liver (96). In both species, we observed that all seven genes associated with this pathway were expressed in both OGM and ODM conditions to varying degrees. Similar patterns of expression were observed between bovine and ovine samples, apart from *G6PC1* where expression was significantly enriched in bovine ODM when compared to OGM. The expression of the genes associated with this pathway represents the conservation of another fundamental liver process within these ruminant organoid cultures and opens the door for the use of this novel platform for the in-depth study of specific metabolic liver processes *in vitro* in ways not previously possible.

Liver organoids have been posited as *in vitro* platforms for screening novel drug compounds and demonstrating drug metabolism and toxicity responses (31,97–101). Here, we demonstrate that bovine and ovine liver organoids not only retain expression of key genes associated with drug metabolism, such as cytochrome P450 and FMO family members, but also display hepatic biotransformation capacity. We demonstrated this activity by modelling the metabolism of triclabendazole (TCBZ), tracking its conversion from the parent compound to its most abundant and pharmacologically active metabolite, TCBZ-SO (102,103). TCBZ was selected as it has a well-defined metabolic pathway and is the primary treatment of livestock and human fascioliasis, an infection of the liver caused by the trematode parasites such as *Fasciola hepatica* and *Fasciola gigantica* (104). Organoids derived from both species could convert TCBZ to TCBZ-SO, with measurable levels detected in both the cellular fraction and the cell culture supernatants. This pattern suggests not only active biotransformation but also secretion of TCBZ-SO by the organoid cultures, further supporting their functional hepatic phenotype. Limitations associated with conducting experimental work in livestock, combined with the absence of suitable *in vitro* models, mean that the precise mechanisms of action of many compounds remain poorly defined. Additionally, toxicity testing often requires costly, large-scale animal trials. The ability of ruminant liver organoids to actively metabolize drug compounds now offers a transformative alternative enabling more efficient initial testing of novel compounds and reducing dependence on *in vivo* trials.

In summary, we report the development of liver organoids containing functional cholangiocytes and hepatocytes from cattle and sheep liver progenitor cells. Comparative analyses with matched tissue samples enables us to delineate distinct immunological and metabolic landscapes of the bovine and ovine livers. The demonstrated functionality of these organoid cultures, including their capacity for active drug metabolism, positions them as valuable new *in vitro* tools for investigating livestock health and disease, while reducing dependency on live animals for exploratory phase research.

## Materials and Methods

### Animals

Bovine liver tissue used in this study was derived from 7 animals from two separate cohorts. The first cohort consisted of four, 8-day old male Holstein-Friesian cross calves and the second consisted of three, 9-month-old male Holstein-Friesian cross calves. Sheep liver tissue was derived from four 9-month-old female, texel cross sheep. This study was carried out in line with the 3Rs principles, specifically regarding the replacement or reduction of animals used in research, and as such all animal tissue used was opportunistically obtained post-mortem from healthy control animals from separate research trials performed at the Moredun Research Institute, UK.

### Isolation of Hepatic Ductal Fragments

Liver tissue was removed from both species at postmortem. Approximately 10 cm^2^ of liver tissue was collected along the common hepatic duct and connecting intrahepatic ducts using a sterile scalpel and forceps, including both the ductal tissue and a thin layer of parenchyma. Tissue was placed into Hank’s buffered saline solution (HBSS) containing 50 *μ*g/ml gentamicin (ThermoFisher Scientific), 100 U/mL penicillin/streptomycin, and 0.25 mg/mL amphotericin B (ThermoFisher Scientific) and placed on ice. Excess parenchymal tissue was then removed, and ductal tissue was minced further using a sterile scalpel into pieces approximately 5 mm^2^ and cryopreserved in Cryostor CS10 (Stem Cell Technologies). Later, resuscitated tissue was placed into a Falcon tube containing 30 mL of HBSS containing 50 µg/mL gentamicin, 100 U/mL penicillin/streptomycin, and 0.25 mg/mL amphotericin B. The tissue was gently shaken, prior to centrifugation at 200 x *g* for 1 minute to allow pieces to settle and supernatant was discarded. For isolation of hepatic ductal fragments containing progenitor cells, tissue digestion media was made using 35 mL advanced Dulbecco’s Modified Eagle Medium/Ham’s F-12 (Adv DMEM/F12 (ThermoFisher Scientific), 4500 U type I collagenase (30130, Sigma-Aldrich), 50 *μ*g/mL gentamicin, 100 U/mL penicillin/streptomycin, and 0.25 mg/ml amphotericin B. The minced tissue was placed into a 50 mL Falcon tube alongside 5 mL of digestion media, shaken gently, and incubated in a 37°C water bath for 15 minutes. The tissue was allowed to settle by gravity, and the supernatant containing hepatic ductal fragments was transferred to a new 15 mL Falcon and centrifuged at 400 *x g* for 5 minutes. Supernatant was removed and the pellet was resuspended in 1 mL Adv DMEM/F12 and placed on ice until all six digests were finished. Five millilitres of digestion media was added to the remaining tissue and incubated for a further 15 minutes prior to repeating this process. Following six rounds of digest and processing of the hepatic ductal fragments, the fragments from each digest were pooled and centrifuged at 400 *x g* for 5 minutes. The pellet was resuspended in 1X ammonium chloride solution for red blood cell lysis and incubated on ice for 5 minutes, prior to centrifugation at 400 *x g* for 5 minutes. This final pellet was resuspended in 1 mL Adv DMEM/F12.

### Organoid Culture

Approximately ¼ of the final isolated hepatic ductal fragments were resuspended in 100 *μ*l of Adv DMEM/F12 and added to 150 *μ*l of Corning® Growth Factor Reduced, Phenol Red-Free, Matrigel™ Basement Matrix (356231; Corning). 50 *μ*l droplets of this mixture were added to the centre of a pre-warmed, 24-well tissue culture plate (CLS3527; Corning) and incubated at 37°C, 5% CO_2_ for 30 minutes to allow the matrix to polymerise. Five hundred microlitres of pre-warmed Hepaticult Organoid Growth Medium (Mouse) (OGM) (StemCell Technologies) containing 500 nM Y-27632 (StemCell Technologies), 5 *μ*M Forskolin (StemCell Technologies), 1X N2 supplement (ThermoFischer), 5 *μ*M A83.01 (StemCell Technologies), and 50 *μ*g/mL gentamicin. For establishment of cultures at passage zero, media was supplemented with 50 *μ*g/mL Primocin (Invivogen). Plates were incubated at 37°C, 5% CO_2_ with Hepaticult medium replaced every 2-3 days. Organoids were kept in culture for 5-7 days prior to passage.

### Organoid Passage and Cryopreservation

Hepaticult medium was removed from each well and replaced with 1 mL of ice-cold Adv DMEM/F12 to dissolve the Matrigel™ domes. The organoids were pooled from each animal into a 15 mL Falcon tube and topped up to 10 mL with Adv DMEM/F12 prior to centrifugation at 300 *x g* for 5 minutes. The supernatant was removed, and organoids were resuspended in 200 *μ*l of Adv DMEM/F12. Organoids were fragmented via mechanical disruption by passing through a 200 *μ*l pipette tip, bent at a 90° angle, approximately 20-30 times. One hundred microlitres of organoid fragments were resuspended in 150 *μ*l of Matrigel™ and seeded into the centre of a 24-well tissue culture plates as previously described.

For cryopreservation of organoids, Hepaticult medium was removed from each well and replaced with 1 mL of ice-cold Adv DMEM/F12. Organoids were then pooled into a 15 mL Falcon tube, topped up to 10 mL with Adv DMEM/F12 and centrifuged at 300 *x g* for 5 minutes. The organoid pellet was then resuspended in CryoStor™ CS10 (StemCell Technologies) at approximately 100-500 organoids per 1 mL CryoStor™ 10 and then 1 mL transferred to each cryovial. Cryovials were placed in a cryogenic freezing container at -70°C overnight, prior to long term storage at -150°C. For resuscitation of cryopreserved organoids, cryovials were placed in a 37°C water bath for 1-2 minutes to rapidly thaw. Organoids were then transferred to a 15 mL Falcon tube containing 9 mL of Adv DMEM/F12 and centrifuged at 350 *x g* for 5 minutes. The organoid pellet was then resuspended in 100 *μ*l of Adv DMEM/F12, mixed with 150 *μ*l of Matrigel™, and seeded into 50 *μ*l domes as described above.

### Organoid Differentiation

Organoids were seeded as described above in Hepaticult OGM and expanded for 3 days. The culture media on day 3 was then changed to Hepaticult Organoid Differentiation Medium (Human; ODM) (StemCell Technologies) containing 500 nM Y-27632 (StemCell Technologies), 3 *μ*M Dexamethasone (StemCell Technologies), 5 *μ*M A83.01, 1X N2 Supplement, and 50 *μ*g/mL gentamicin. For ovine differentiation, Hepaticult ODM was also supplemented with 0.1 *μ*g/mL recombinant human Tumour Necrosis Factor-α (TNF-α; PHC3015, Gibco). Organoids were sustained in differentiation medium for a further 5-7 days and observed via light and phase contrast microscopy.

Where we sought to determine whether modifications to TNF-α and A83.01 would influence differentiation in ovine organoids, Hepaticult ODM was supplemented with and without 5 *μ*M A83.01 as well as with and without varying concentrations of TNF-α (0.1 µg/mL and 0.5 µg/mL). Hepaticult ODM was further supplemented with 500 nM Y-27632, 1X N2 supplement, 3 *μ*M Dexamethasone, and 50 *μ*g/mL Gentamicin.

### Immunohistochemistry

For immunohistochemical (IHC) analysis of tissue, samples were collected at postmortem from sites immediately adjacent to where tissue was harvested for organoid generation and fixed for 24h at room temperature (RT) in 10% neutral buffer formalin prior to storage at 4°C in 70% ethanol. For IHC analysis of organoids, organoids were grown in either growth or differentiation conditions as described above. Hepaticult medium was removed from each well and replaced with 1 mL of ice cold Adv DMEM/F12. Three to five wells of organoids per animal were pooled in a 15 mL Falcon topped up to 10 mL with Adv DMEM/F12 and centrifuged at 300 *x g* for 5 minutes. Once removed from the Matrigel organoids needed to be handled gently as they were fragile and can break apart. For fixation, organoids were resuspended in 1 mL of 10% neutral buffer formalin and left at RT for 20 minutes, prior to centrifugation at 300 *x g* for 5 minutes. Fixed organoids were washed twice with phosphate buffer saline (PBS) and then resuspended in 100 *μ*l of Epredia™ HistoGel™ Specimen Processing Gel and placed into a disposable histology base mould (Fischer Scientific) on ice for 15 minutes. The Histogel™ block containing the organoids was then transferred to a histology cassette and stored at 4°C in PBS prior to embedding. The fixed organoids (in Histogel) and tissues were embedded in paraffin blocks. Four micrometre thick sections were cut, placed on Epredia™ SuperFrost Plus™ Adhesion slides (Fischer Scientific), dried overnight at 37°C and stored long-term at 4°C.

Prior to fluorescence antibody labelling, sections underwent xylene- and ethanol-based deparaffinisation prior to antigen retrieval in citrate buffer for 15 minutes at 95°C. Slides were permeabilised for 10 minutes using 0.1 % Triton-X 100 (ThermoFisher Scientific) in PBS. Following permeabilization, slides were washed twice using PBST (PBS containing 0.5% Tween 20 (ThermoFisher Scientific) prior to blocking with 3% cold water fish gelatin (ThermoFisher Scientific) dissolved in PBST. Slides were washed once in PBST prior to addition of the primary antibodies for overnight incubation at 4°C. A full list of primary and secondary antibodies used in this study is found in Table 1. Following overnight incubation, slides were washed three times with PBST prior to incubation with the designated secondary antibody for 1 hour at RT. Slides were washed three times with PBST, counterstained for 20 minutes with Hoechst 33258 nuclear dye (94403, Sigma-Aldrich), washed an additional three times in PBST and mounted using ProLong™ Diamond Antifade Mountant (ThermoFischer Scientific). Slides were imaged using a Zeiss Axio Observer 7 and processed using ZenBlue (v. 3.4.91). Slides were stored for up to one month at 4°C.

**Table 1.** Primary and secondary antibodies used in this study.

| Target | Host | Working Dilution | Product Information |
| --- | --- | --- | --- |
| KRT19 | Mouse | 1:300 | SC376126, Santa Cruz Biotechnology |
| Bovine Serum Albumin (BSA) | Rabbit | 1:300 | A11133, ThermoFisher Scientific |
| Ki67 | Rabbit | 1:500 | Ab15580, Abcam |
| KRT18 | Mouse | 1:300 | Ab668, Abcam |
| Hoechst 33258 |  | 1:1000 | 94403, Sigma-Aldrich |
| Rabbit IgG AF488 | Goat | 1:500 | A11008, ThermoFisher Scientific |
| Mouse IgG AF488 | Goat | 1:500 | A11001, ThermoFisher Scientific |

### Total RNA Extraction

Liver tissue was collected from sites immediately adjacent to where material was harvested for organoid culture. Total RNA was extracted from bovine and ovine whole liver tissue stored in RNA*Later* solution (ThermoFischer), bovine organoids in growth and differentiation medium at passage 1 or 2, and ovine organoids in growth medium at passage 2 and 7 and differentiation medium at passage 2. Organoids in both growth and differentiation conditions were cultured as described above, with 5 wells per condition being pooled for total RNA extractions. OGM-cultured organoids were maintained for 7 days prior to collection. For total RNA extraction, Hepaticult medium was removed and 1 mL of ice-cold Adv DMEM/F12 was added to each well to dissolve the Matrigel™ matrix. Organoids were transferred to a 15 mL Falcon tube and topped up to 10 mL with Adv DMEM/F12 prior to centrifugation at 300 *x g* for 5 minutes. Supernatants were removed and the organoid pellets were resuspended in 350 *μ*l buffer RLT (Qiagen) containing 3.5 *μ*l β-mercaptoethanol (β-ME) and immediately stored at -150°C. The addition of β-ME to organoids and tissue was essential to inhibit RNase activity and retain RNA integrity, due to the high levels of RNase associated with liver material. Total RNA extraction was performed using the RNeasy Mini Kit (Qiagen) with the optional on-column DNase digest according to manufacturer’s instructions, with final total RNA elution of 30 *μ*l in nuclease-free water. Total RNA was quantified using a Nanodrop™ and RNA integrity was tested using a Bioanalyzer (Aligent) and the total RNA 6000 Nano kit. All samples in this study obtained a bioanalyzer score of 6.5 or above. Total RNA was stored at -70°C until RNA sequencing.

For bovine and ovine liver total RNA extraction, approximately 30 mg of liver tissue previously collected at postmortem and stored in RNA*later* solution (ThermoFischer) was homogenized in 600 *μ*l RLT buffer containing 6 *μ*l β-ME using a Precellys® Tissue Homogeniser with ZR BashingBead™ Lysis Tubes (0.1 and 2.0 mm) (Cambridge Bioscience). Tissue homogenisation occurred using 2 x 15 second, 6500 rpm pulses. Samples were centrifuged at maximum speed for 1 minute to remove residual cell debris. Total RNA extraction utilised the RNeasy Mini Kit according to manufacturer’s instructions, with final elution of 30 *μ*l in nuclease free water. RNA quantity and integrity was tested as described above and final total RNA samples were stored at -70° C.

### RNA-Sequencing Analysis

All library preparation and sequencing were conducted at Novogene (Cambridge, UK), utilising the Illumina PE150 Novoseq platform for eukaryotic mRNA sequencing with Poly(A) enrichment for paired end 150 base pair (bp) reads and a 30 million sequencing read depth. A total of 37 libraries were constructed [including: ovine growth organoids P2 and P7; ovine differentiated organoids P2; ovine liver tissues (n = 4); bovine growth organoids P1/2; bovine differentiated organoids P1/2; bovine liver tissue (n = 7; 3 and 4 from two cohorts, respectively)]. Sequenced files were checked for quality using FastQC (v 0.12.1). Sequenced reads were pseudo-aligned to either the *Ovis aries* (ARS-UI_Ramb_v2.0) or *Bos taurus* (ARS-UCD1.3) genomes using Kallisto (v 0.51.1) (70-75% genome mapping rates across every sample). For comparison between bovine and ovine samples, direct gene orthologues were identified using biomart in RStudio (v.024.12.1+563). Differential gene expression analysis between samples was performed using DESeq2 (v1.46.0) with the transcript-level quantification (abundance) files produced from Kallisto pseudo-alignment (105). Resulting genes were filtered to include only those with padj > 0.05, log_2_ fold change > 1, and a normalized count > 10. Transcript abundance calculated as transcripts per million (TPM) were also generated from Kallisto pseudo-alignment and used for gene comparisons within an individual sample.

For cluster enrichment analysis, liver cell clusters were identified from Aizarain et. al. (2019) and filtered to include only genes with a log_2_ fold change >2 for each cluster. These clusters were then compared to the differentially expressed genes identified through DESeq2 analysis for comparisons between tissue versus growth organoids and differentiated versus growth organoids for both species, respectively. RStudio (version 024.12.1+563) was used for all analysis of abundance files following Kallisto pseudo-alignment. Gene set enrichment analysis was performed using RStudio package fgsea (version 1.32.4). Enrichment was tested using fgsea multilevel with gene set size constraints (min size = 10, max size = 500) and 10,000 simple permutations for initial p-value estimation where specified. Normalized enrichment scores (NES) and FDR-adjusted p-values were reported. Enrichment curves were visualized using fgsea’s plot enrichment. PCA plots were generated using prcomp and all plots were generated using ggplot2 (v. 4.0.1).

### RT-qPCR

Ovine liver organoids were grown in Hepaticult OGM for three days prior to differentiation. Following 5 days in culture, RNA was extracted as previously described, quantified using a Nanodrop™, and stored at -70° C prior to cDNA synthesis. SuperScript™ III Reverse Transcriptase (ThermoFisher Scientific) was used for cDNA synthesis, alongside RNaseOUT Recombinant Ribonuclease Inhibitor (ThermoFisher Scientific), and Ambion™ RNase H (ThermoFisher Scientific) following manufacturer’s instructions. For RT-qPCR reactions, PowerTrack™ SYBR Green Master Mix was added to 1 *μ*l synthesized cDNA, 0.2 µM forward and reverse primers for *ALB*, *GAPDH*, and *ACTB* per reaction. Primer sequences were designed as shown in Table 2. PCR reactions were performed on an Applied Biosystems 7500 Real Time PCR System (Applied Biosystems, Foster City, CA, USA). Thermal cycling conditions were 94 °C for 2 min; followed 40 cycles of 94 °C for 15 s, 60 °C for 30 s and 72 °C for 30 s. For qualitative analysis of *ALB* gene expression, Δct values were normalised to the housekeeping genes *GAPDH* and *ACTB*. The ΔΔt values were normalised to the Δct values of organoids grown in OGM.

**Table 2.**
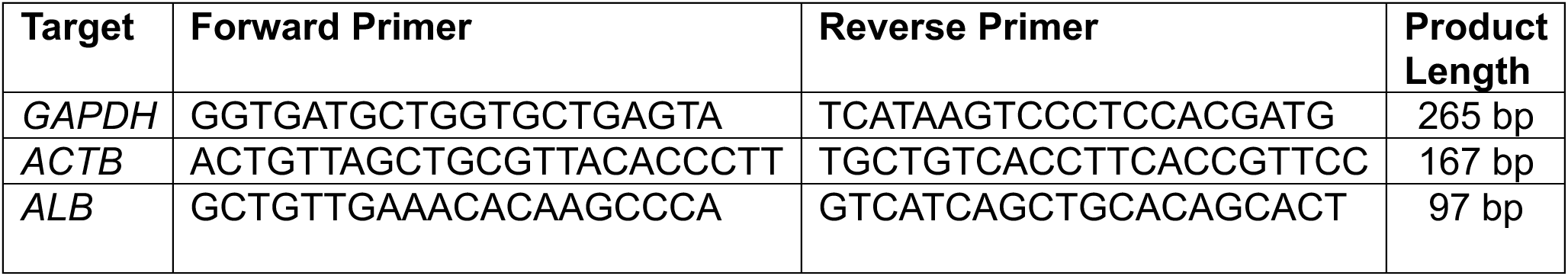
Primer sequences used for RT-qPCR.

### Determination of TCBZ and its metabolites in liver organoids by LC-MS/MS

Bovine and ovine liver organoids, cultured in either standard (OGM) or differentiation (ODM) conditions, were treated with 20 µM TCBZ (Sigma-Aldrich, UK) in their respective media condition for 24 h and extracted according to Cai et al. (2010). After the incubation period, the cell culture supernatants were collected and immediately frozen at -70 °C prior to analysis. The organoids were removed from Matrigel™ as described previously and washed once with ice cold Advanced DMEM to remove remaining extracellular matrix. The organoids were pelleted via centrifugation at 400 x *g* for 5 minutes and frozen at -70 °C prior to extraction. For metabolite extraction, each organoid pellet was resuspended in 1 mL of acetonitrile (ACN) and vortexed briefly. Residual lipids were removed from the samples via hexane extraction, whereas 1 mL of hexane (ThermoFisher Scientific) was added to each sample and vortexed for 30 seconds. The upper hexane phase containing excess lipids was then removed from each sample and this process was repeated three times. Following the final round of hexane extraction, the samples were frozen at -70 °C prior to analysis.

TCBZ, TCBZ-SO, TCBZ-SO_2_ and keto-TCBZ (Sigma-Aldrich, UK) standards were prepared in methanol and used to construct a six-point internal calibration curve (0-500 µg/L for TCBZ and 0-1000 µg/L for the metabolites). Peak areas were used as the analytical response versus concentration. For chromatographic analysis, 5-µl samples were injected into an Agilent 1200 LC chromatography system equipped with an Acquity HSS T3 column (2.1 mm x 100 mm, particle size 1.8 µM; Waters, ROI) at a flow rate of 0.5 ml/min-1. Mass spectrometric detection was performed using an Agilent 6495B Triple Quadrupole mass spectrometer in dynamic MRM mode. The following targeted mass transitions were used: for Keto-TCBZ 329 m/z (precursor ion) and 168 m/z (product ion), for TCBZ 359 m/z (precursor ion) and 344 m/z (production), for TCBZ-SO 375 m/z (precursor ion) and 357 m/z (product ion) and for TCBZ-SO2 391 m/z (precursor ion) and 241 m/z (product ion). Agilent MassHunter (Version 10.1) was used for data acquisition whilst data processing was performed with Agilent Quantitative Analysis software (Version 10.1). The full LC-MS/MS instrument parameters are shown in Supplementary Table 1.

## Supporting Information

Supplemental Figure 1: Bovine Heatmaps

Supplemental Figure 2: Ovine Heatmaps

Supplemental Figure 3: ALB Tissue Labelling

Supplemental Figure 4: KRT19 ODM

Supplemental Figure 5: GO Analysis

Supplemental Figure 6: Ovine Diff TNF-a

Supplemental File 1: Human Cluster

Supplemental File 2: Bovine Normalized Counts and DEGs

Supplemental File 3: Ovine Normalised Counts and DEGs

Supplemental File 4: Species Comparison DEGs

Supplemental File 5: LC-MS/MS TCBZ Raw Data

Supplemental Table 1: MS Settings for TCBZ

## Acknowledgements

We would like to thank the Bioservices Unit at the Moredun Research Institute, UK for their assistance in the provision of liver tissue at postmortem. We would also like to thank Dr Dermot Faulkner (Agri-Food & Biosciences Institute) for providing TCBZ reference standards for LC-MS analysis.

