## Supplemental Figures for "A comparative analysis of liver tissue and novel primary organoid cultures from ruminants reveals species-specific immune architecture and metabolic specialization"

### Slide 1
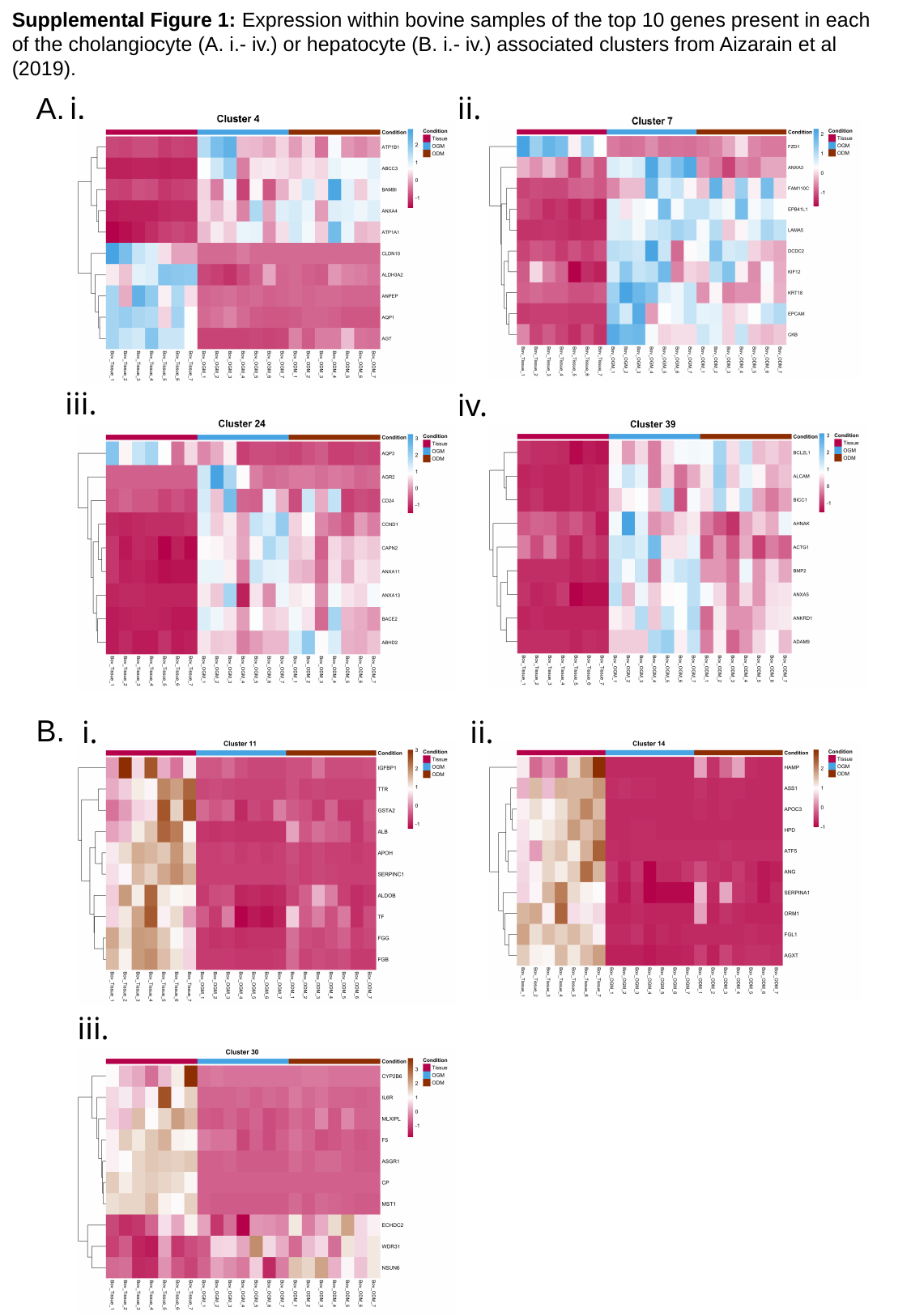

Supplemental Figure 1: Expression within bovine samples of the top 10 genes present in each of the cholangiocyte (A. i.- iv.) or hepatocyte (B. i.- iv.) associated clusters from Aizarain et al (2019).
A.
i.
ii.
iii.
iv.
B.
i.
ii.
iii.

### Slide 2
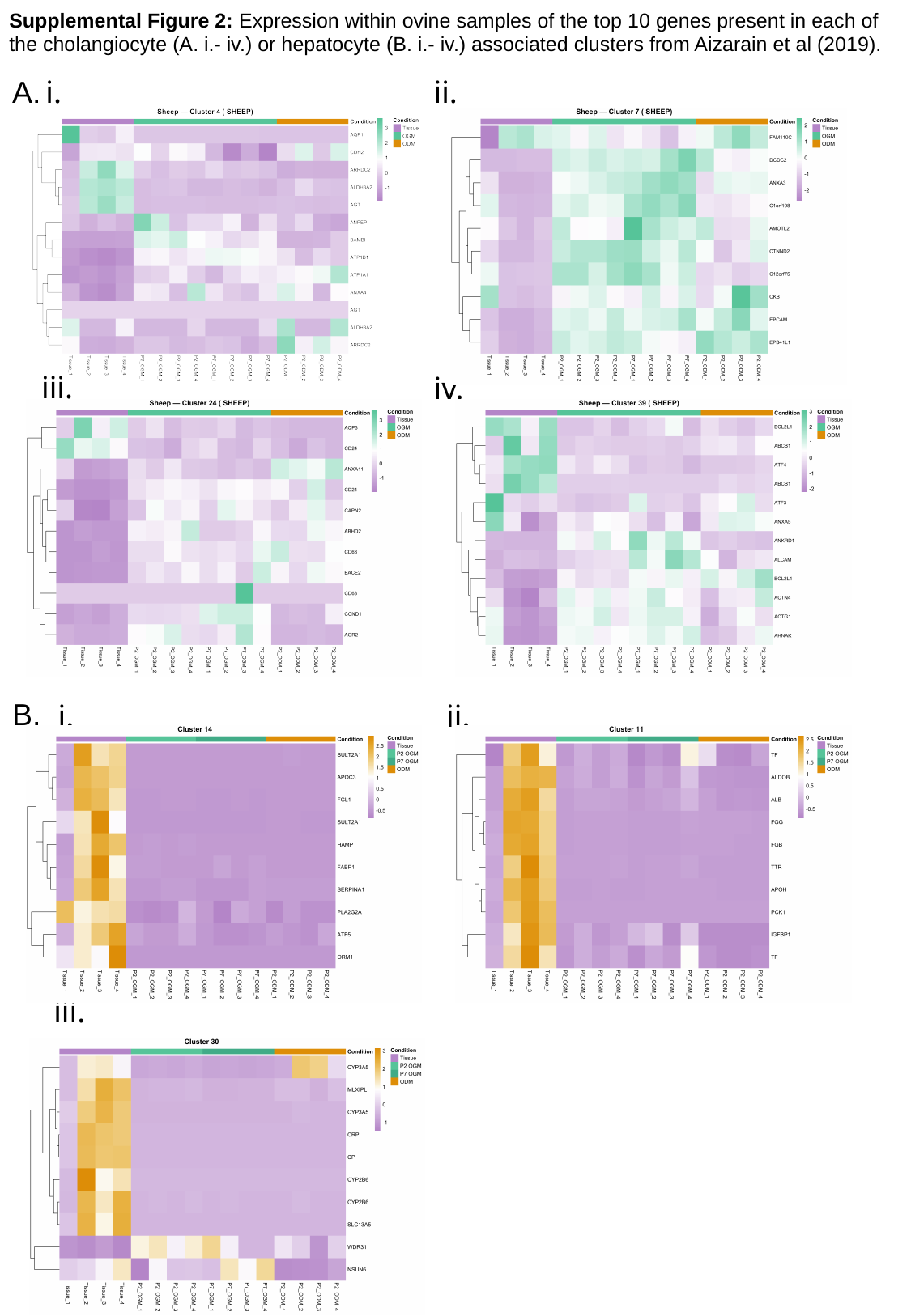

Supplemental Figure 2: Expression within ovine samples of the top 10 genes present in each of the cholangiocyte (A. i.- iv.) or hepatocyte (B. i.- iv.) associated clusters from Aizarain et al (2019).
A.
i.
ii.
iii.
iv.
B.
i.
ii.
iii.

### Slide 3
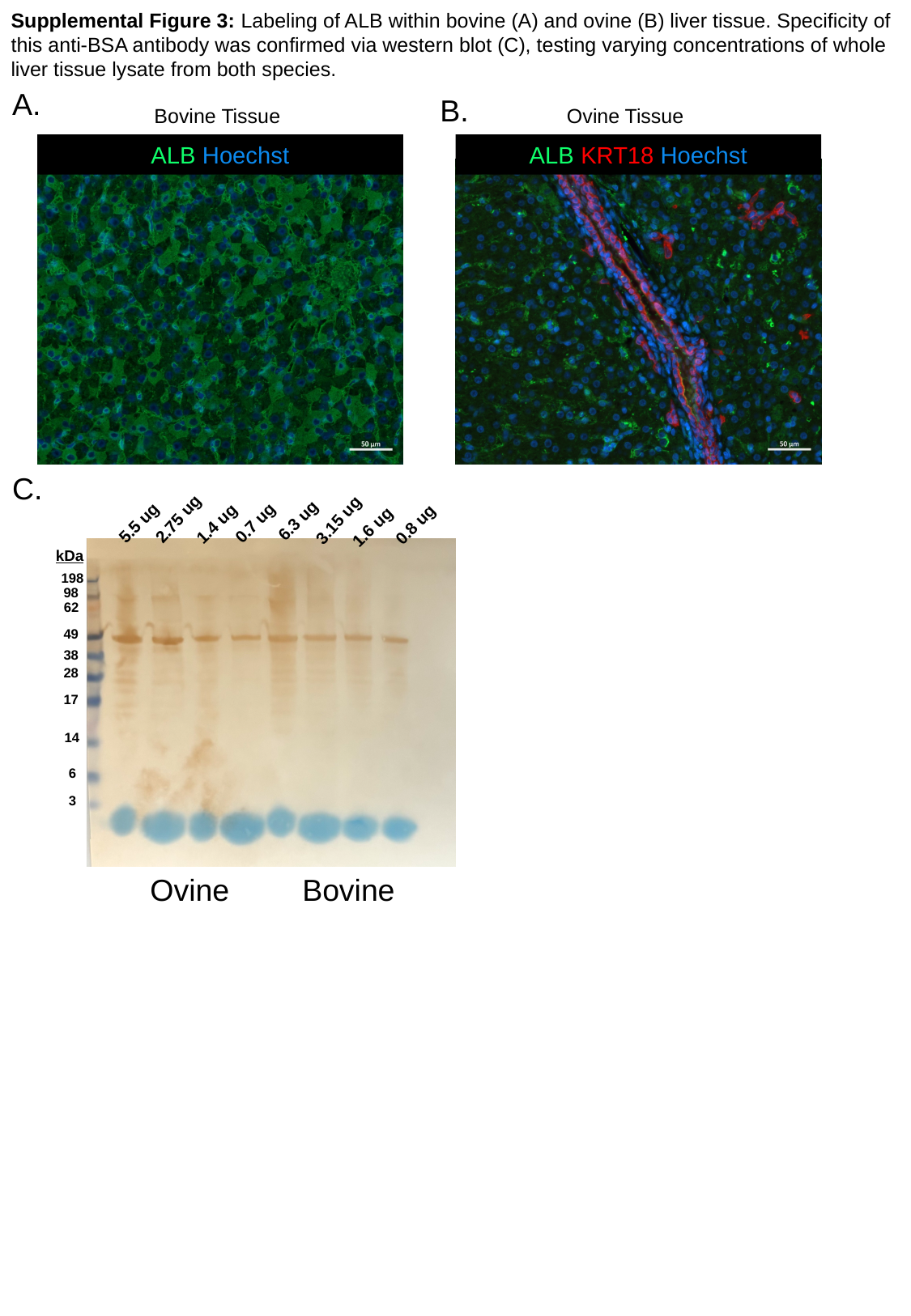

Supplemental Figure 3: Labeling of ALB within bovine (A) and ovine (B) liver tissue. Specificity of this anti-BSA antibody was confirmed via western blot (C), testing varying concentrations of whole liver tissue lysate from both species.
A.
B.
Ovine Tissue
Bovine Tissue
ALB Hoechst
ALB KRT18 Hoechst
0.7 ug
0.8 ug
1.4 ug
2.75 ug
1.6 ug
3.15 ug
6.3 ug
5.5 ug
Ovine
Bovine
kDa
198
62
49
38
28
17
14
6
3
98
C.

### Slide 4
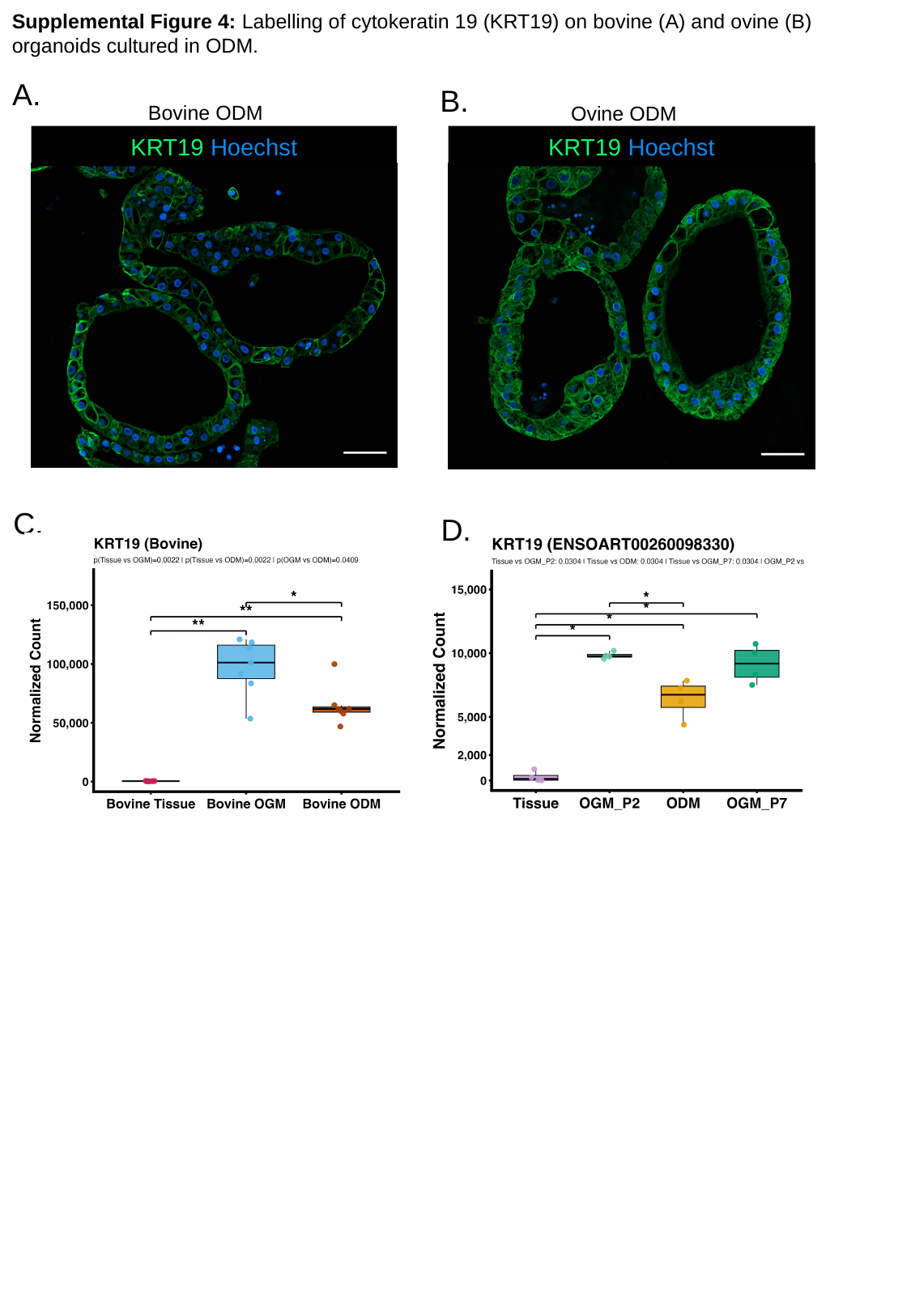

Supplemental Figure 4: Labelling of cytokeratin 19 (KRT19) on bovine (A) and ovine (B) organoids cultured in ODM.
A.
B.
Bovine ODM
Ovine ODM
KRT19 Hoechst
KRT19 Hoechst
C.
D.

### Slide 5
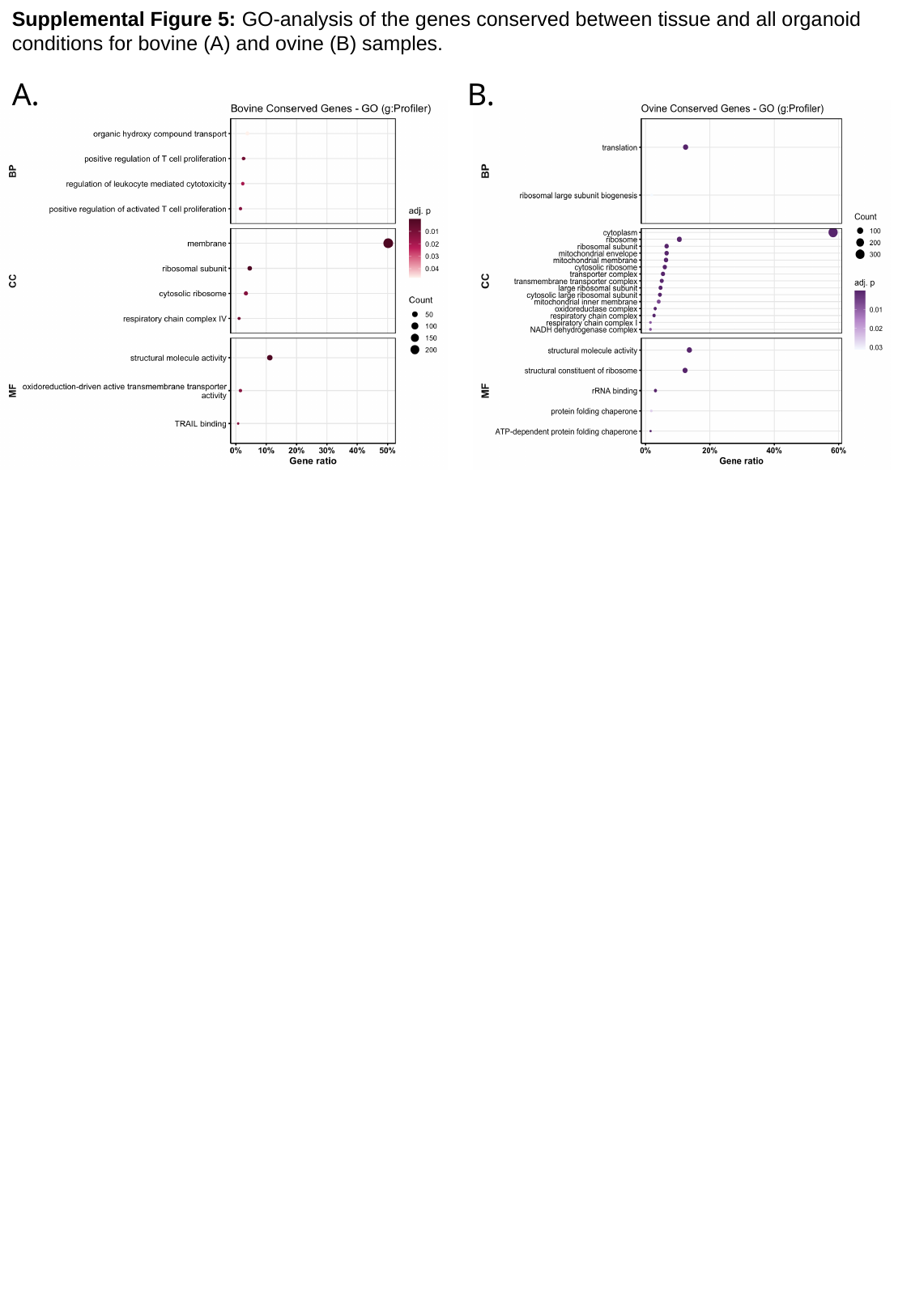

Supplemental Figure 5: GO-analysis of the genes conserved between tissue and all organoid conditions for bovine (A) and ovine (B) samples.
A.
B.

### Slide 6
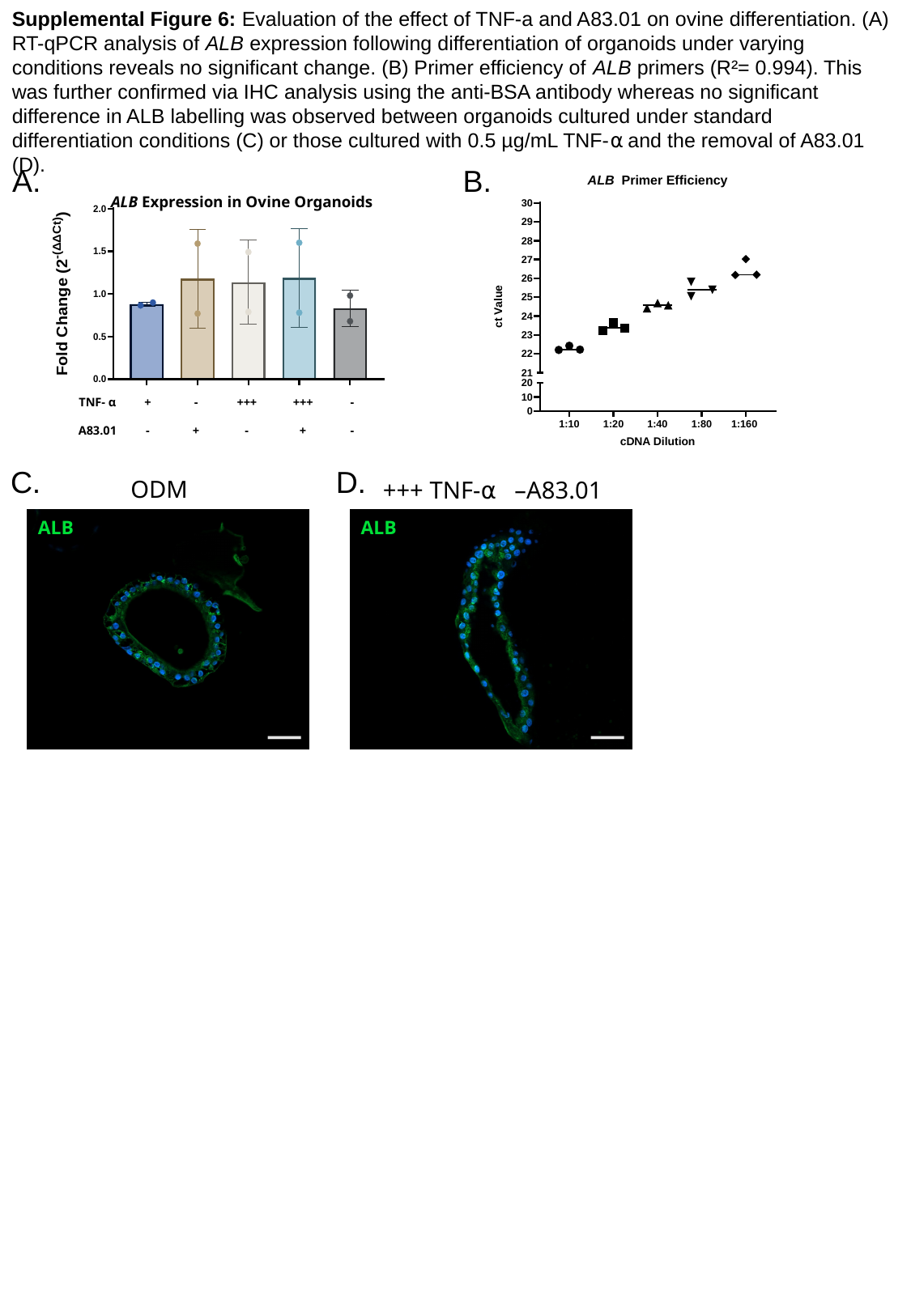

Supplemental Figure 6: Evaluation of the effect of TNF-a and A83.01 on ovine differentiation. (A) RT-qPCR analysis of ALB expression following differentiation of organoids under varying conditions reveals no significant change. (B) Primer efficiency of ALB primers (R²= 0.994). This was further confirmed via IHC analysis using the anti-BSA antibody whereas no significant difference in ALB labelling was observed between organoids cultured under standard differentiation conditions (C) or those cultured with 0.5 µg/mL TNF-⍺ and the removal of A83.01 (D).
A.
B.
ALB Expression in Ovine Organoids
| TNF- α | + | - | +++ | +++ | - |
| --- | --- | --- | --- | --- | --- |
| A83.01 | - | + | - | + | - |
C.
D.
ODM
+++ TNF-⍺ –A83.01
ALB
ALB
