## Supplemental Table 1 for "A comparative analysis of liver tissue and novel primary organoid cultures from ruminants reveals species-specific immune architecture and metabolic specialization"

| LC | Solvents | Solvent A:  Water (0.1% Formic acid)  Solvent B:  Acetonitrile (0.1% Formic acid) |
| --- | --- | --- |
|  | Gradient | *Time (min) %A %B*  0 95 5  14 5 95  15 95 5 |
|  | Column | Type: HSS T3  (Waters Acquity 186003539)  Pore size: 100Å  Particle size: 1.8 μm  2.1 mm x 100 mm |
|  | Column Temperature | 30 C° |
|  | Sample Volume | 5 μl |
|  | Flow | 0.5 ml min^-1^ |
| MS | Acquisition Mode | Dynamic MRM |
|  | Polarity | Positive |
|  | Fragmentor Voltage | 300 V |
|  | Ion Source | AJS ESI |
|  | Accelerator Voltage | 5V |
|  | Collision Energy (V) | Keto-TCBZ: 30  TCBZ: 30  TCBZ-SO: 22  TCBZ-SO_2_: 30 |

**Supplementary Table 1**: Parameters for chromatographic separation (LC) and detection (MS) of TCBZ metabolites using an Agilent LC-MS/MS 6495B.
